# A chimeric Japanese encephalitis vaccine protects against lethal yellow fever virus infection without inducing neutralizing antibodies

**DOI:** 10.1101/717181

**Authors:** Niraj Mishra, Robbert Boudewijns, Michael A. Schmid, Rafael Elias Marques, Sapna Sharma, Johan Neyts, Kai Dallmeier

**Author notes:** **<u>Correspondence: co-corresponding</u> authors**, Johan Neyts and Kai Dallmeier <u>ADDRESS FOR CORRESPONDENCE:</u> KU Leuven Department of Microbiology and Immunology, Rega Institute, Laboratory of Virology and Chemotherapy, Herestraat 49 (postbus 1043), B-3000 Leuven, Belgium.

## Abstract

Recent massive outbreaks of yellow fever virus (YFV) in West Africa and Brazil resulted in rapid depletion of global vaccine emergency stockpiles and raised concerns about being not prepared against future YFV epidemics. Here we report that a live-attenuated virus similar to the Japanese encephalitis virus (JEV) vaccine JE-CVax/Imojev^®^ that consists of YFV-17D vaccine from which the structural (prM/E) genes have been replaced with those of the JEV SA14-14-2 vaccine strain confers full protection in mice against lethal YFV challenge. In contrast to the YFV-17D mediated protection against YFV, this protection is not mediated by neutralizing antibodies but correlates with YFV-specific non-neutralizing antibodies and T cell responses against cell-associated YFV NS1 and other YFV non-structural (NS) proteins. Our findings reveal the importance of YFV NS proteins to mediate protection and demonstrate that chimeric flavivirus vaccines, such as Imojev^®^ can confer protection against two flaviviruses. This dual protection has implications for the possible off-label use of JE-CVax in case of emergency and vaccine shortage during YFV outbreaks. In addition, populations in Asia that have been vaccinated with Imojev^®^ may already be protected against YFV should outbreaks ever occur on that continent as feared by WHO.

**IMPORTANCE:** Efficient and safe vaccines exist against yellow fever (e.g. YFV-17D) that provide long-lasting protection by rapidly inducing neutralizing antibody responses. However, vaccine supply cannot cope with an increasing demand posed by massive urban outbreaks in recent years. Here we report that JE-CVax/Imojev^®^, a YFV-17D-based chimeric Japanese encephalitis vaccine also efficiently protects against YFV infection in mice. In case of shortage of the YFV vaccine during yellow fever outbreaks, (off-label) use of JE-CVax/Imojev^®^ may be considered. Moreover, wider use of JE-CVax/Imojev^®^ in Asia may lower the risk of the much-feared YFV spill over to the continent. More in general chimeric vaccines that combine surface antigens and replication machineries of two distinct flaviviruses can be considered dual vaccines, for the latter pathogen without induction of surface-specific antibodies. Following this rationale, novel flavivirus vaccines that do not hold a risk for antibody-dependent enhancement (ADE) of infection [inherent to current dengue vaccines and dengue vaccine candidates] could be designed.

---

Several flaviviruses, such as the yellow fever virus (YFV), Japanese encephalitis virus (JEV), dengue virus (DENV), Zika virus (ZIKV), West Nile virus (WNV) and tick-borne encephalitis virus are important human pathogens. Flaviviruses are spread worldwide, though some species show a pronounced restriction to defined endemic regions, such as YFV to sub-Saharan Africa and tropical Latin America, or JEV to Southeast Asia and the Asian Pacific. Certain flaviviruses such as DENV, WNV and most recently ZIKV are (re-)emerging in new areas (1–3). Some evidence suggests the first autochthonous transmission of JEV in Africa (4).

Yellow fever (YF) is an acute viral haemorrhagic disease, which is currently endemic to ∼50 countries with ∼1 billion people living at risk of infection. Despite the availability of a highly efficient vaccine (YFV-17D; e.g. Stamaril^®^), an estimated ∼0.2 million YFV infections with 29,000-60,000 deaths occur annually (5). Recent YFV outbreaks in Angola (2015/16), the Democratic Republic of the Congo (2016), Brazil (2017), and Nigeria (2018), and shortage of the YF vaccine supply raised serious concerns about the preparedness for future outbreaks (6, 7). Since, the *Aedes aegypti* mosquito, the main YFV vector, is omnipresent in (sub)tropical Asia, YFV spill over to Asia and the establishment of epidemics involving urban transmission becomes increasingly realistic (8, 9). For JEV, two licensed vaccines are available, namely Ixiaro^®^ (inactivated vaccine) and JE-CVax (Imojev^®^: YFV-17D-based chimeric live-attenuated vaccine; c-LAV) (10, 11). Another YFV-17D-based tetravalent c-LAV, namely against dengue (CYD-TDV, Dengvaxia^®^) has reached marketing licensure and is being introduced in some countries/regions. However, there are serious concerns related to the use of this vaccine, mainly because of aggravation of dengue disease by pre-existing antibodies (antibody-dependent enhancement, ADE) of DENV infection (12–14).

Vaccination against flaviviruses generally relies on the strategy to mount protective humoral immunity against structural proteins, in particular neutralizing antibodies (nAbs) elicited against the viral envelope (E) protein (5, 15); though also CD4^+^ T cells seem to contribute to the protective activity of current YFV vaccines (16). Nonetheless, experimental evidence in mice and non-human primates for YFV (17–20), and more recently in mice also for WNV (21) and ZIKV (22) clearly shows that also non-structural (NS) proteins, in particular NS1, can evoke protective humoral and cellular immune responses. Of note, NS1 is not part of the infectious flavivirus particle and thus not target of nAbs. Likewise, immunization with an adenovirus-vector encoding the NS3 protein of YFV-17D elicited strong CD8^+^ T cell responses, which resulted in some degree of protection in mice against subsequent challenge (23). However, full protection was only observed when the vaccine included the structural proteins of YFV-17D as antigen as well (23, 24), obviously in line with the accepted role nAb play in YFV infection. Thus, besides humoral immune responses against the E protein, cellular immune responses against the NS proteins may to some extent also contribute to the immunity against flaviviruses. However, no flavivirus vaccines have been developed nor licensed yet for human use that are based on any of these NS proteins as target antigen. Intriguingly, the genome of chimeric flavivirus vaccines (JE-CVax/Imojev^®^ or CYD-TDV/Dengvaxia^®^) consists of sequences of antigenically distinct flaviviruses (respectively JEV and YFV-17D, and DENV and YFV-17D) and may therefore exert some dual protective activities. Here we demonstrate that vaccination of mice with a construct similar to JE-CVax/Imojev^®^ provides rapidly complete protection against a massively lethal YFV challenge, with a single dose being sufficient for full efficacy. Moreover, we show that this protection is, albeit its unexpected potency, not mediated by nAbs, but by multiple complementary and vigorous responses directed against the NS proteins of YFV-17D.

## RESULTS

### JE-CVax provides full dual protection against lethal JEV and YFV challenge in mice

JE-CVax is a c-LAV that consists of the YFV-17D genome, of which the prM and E genes have been replaced by the corresponding sequences of JEV SA14-14-2. AG129 mice were vaccinated with either 10^3^ or 10^4^ PFU of JE-CVax and 28 days later challenged with 10^3^ PFU [equivalent of 1000 LD_50_] of YFV. This resulted in, respectively, 80 and 100% survival, while YFV infection was uniformly lethal in all non-vaccinated controls (Fig. S1A). Therefore, throughout the further study animals were vaccinated and challenged with 10^4^ PFU JE-CVax (full survival in vaccinated mice) and 10^3^ PFU of YFV (full mortality in non-vaccinated mice), respectively. JE-CVax has originally been developed as a JEV vaccine. As expected, unlike non-vaccinated animals (n = 16), all AG129 mice vaccinated with either JE-CVax (10^2^, 10^3^ or 10^4^ PFU; n ≤ 6;) or the inactivated JEV vaccine Ixiaro^®^ (n=10, 2 times × 1μg: twice 1/6^th^ human dose) (25) were completely protected (p-value; >0.0001) against lethal JEV challenge (Fig. 1A). Remarkably, vaccination with JE-CVax resulted also in 97% survival (n=35/36) against a massively lethal YFV challenge (Fig. 1B). All placebo- (n=38) or Ixiaro^®^- vaccinated (n=12) animals had to be euthanized for humane reasons (mean day to euthanasia [MDE]; 14.6 ± 2.8 days and 15.4 ± 3.5, p-value; >0.0001). Importantly, JE-CVax also conferred similarly vigorous protection against YFV in C57BL/6 wild-type (wt) mice (n=16) against intracranial (i.c.) challenge with 10^4^ PFU of YFV (Fig. 1D). In AG129 mice, a benefit (60% survival) could already be observed 7 days post-vaccination (dpv). At 14 dpv or later all animals were fully protected against lethal challenge (Fig. 1C). To establish that JE-CVax-mediated protection against YFV is specific and not resulting from some residual cross-reactivity as previously observed for certain flaviviruses in mice (26), we challenged age-matched non-vaccinated (n=7), or JE-CVax-vaccinated and YFV-17D-challenged (n=6) AG129 mice with 10^4^ PFU of the more distantly related ZIKV (strain MR766). No protective activity was observed (MDE: non-vaccinated *vs.* vaccinated mice; 23.5 ± 5.4 and 17.4 ± 8.8 days, p-value: 0.4831) (Fig. S1B). Thus, a single-dose immunization with JE-CVax provides a fast (≤14 dpv) and virus-specific protection against lethal YFV exposure in mice.

**FIG 1.**
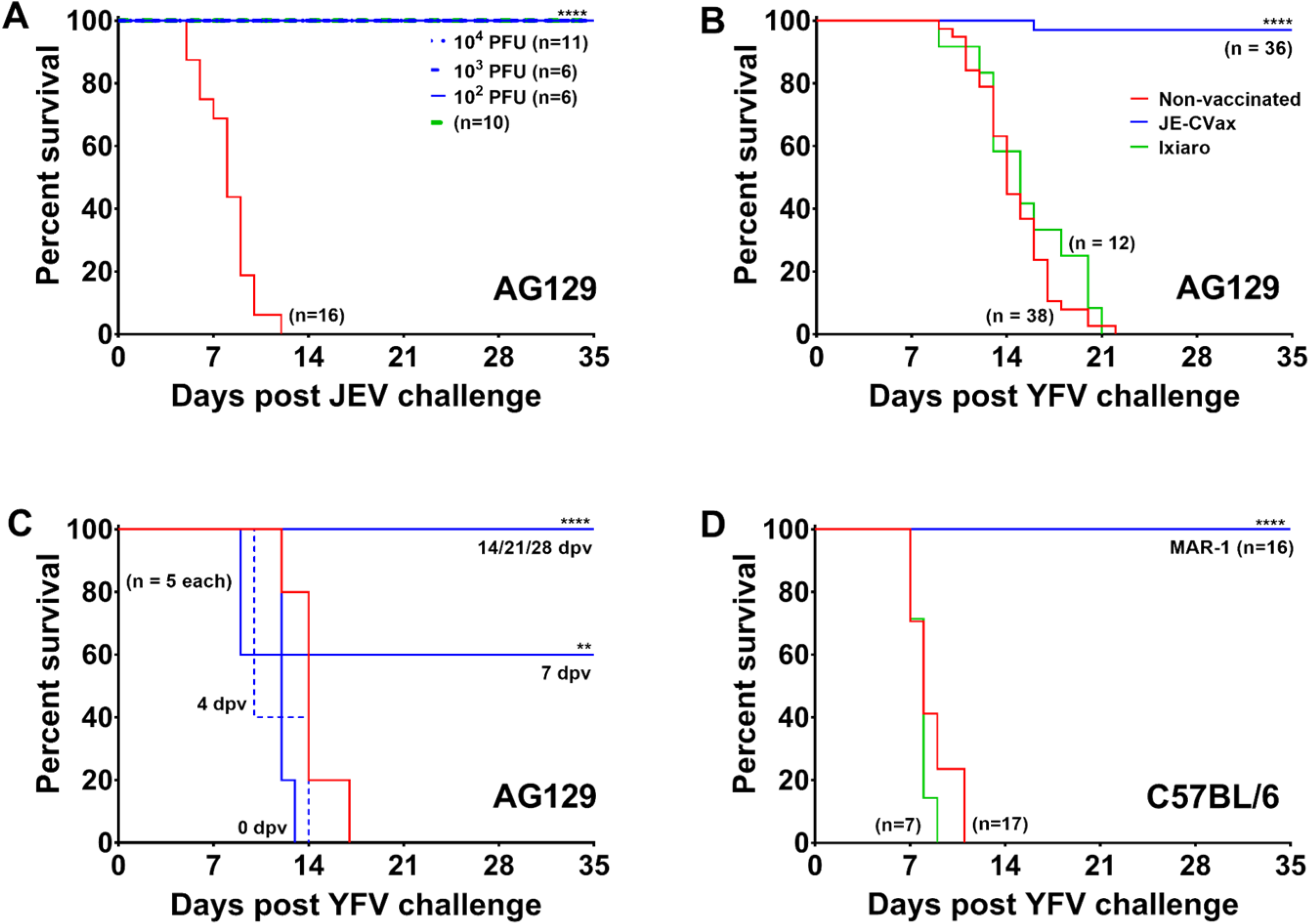
*In vivo* evaluation of JE-CVax-mediated dual protection against lethal JEV SA14-14-2 and YFV-17D challenge. **(A-D)** AG129 and C57BL/6 mice were first vaccinated via the i.p. route with either 10^4^ PFU of JE-CVax (blue), 1 /6th of a human dose of Ixiaro^®^ (green), or assay medium as negative control (red). Animals vaccinated with Ixiaro^®^ were boosted with another 1/6th of a human dose of Ixiaro^®^ 14 dpv. In order to facilitate vaccine virus replication (28), wild-type C57BL/6 mice receiving JE-CVax vaccination were treated with MAR1-5A3 antibody. AG129 mice were i.p. challenged with either 10^3^ of PFU JEV SA14-14-2 at 28 dpv **(A)**, or with 10^3^ PFU of YFV-17D at **(B)** 28 dpv, or **(C)** at 0, 4, 7, 14, 21 and 28 dpv. C57BL/6 mice were i.c. challenged with 10^4^ PFU of YFV-17D at 28 dpv **(D)**. Animals were observed for five weeks after challenge and were euthanized when humane endpoints were reached. The data (A-C) shown represent cumulative results of at least two independent experiments. Log-rank (Mantel-Cox) survival analysis test was performed for statistical significance. ** p-value ≤ 0.01 and **** p-value < 0.001 compared to the non-vaccinated group.

### JE-CVax mediates protection against YFV without involvement of nAbs

To explore whether humoral immunity is involved in JE-CVax-mediated protection against YFV, serum of AG129 mice (i) at day 0 (pre-vaccinated), (ii) infected with YFV-17D before euthanasia (terminal serum), (iii) vaccinated with JE-CVax (day 28; post-vaccinated) or (iv) vaccinated with JE-CVax and challenged with YFV-17D (day 56; post-challenge) was analysed for total binding antibodies and nAb. All animals vaccinated with JE-CVax or Ixiaro^®^ seroconverted to JEV. Sera of non-vaccinated animals that had been infected with YFV-17D showed only some residual reactivity for JEV (as detected by indirect immune fluorescence assay, IIFA; Fig. S2). By contrast, nAbs against JEV were exclusively detected in serum samples of JE-CVax- or Ixiaro^®^-vaccinated animals (log_10_CPENT_50_: 2.48 ± 0.29 or 1.86 ± 0.36, respectively) (Fig. 2A). Only when JE-CVax-vaccinated mice were challenged at a later stage with YFV-17D, nAbs against the latter virus were raised (log_10_CPENT_50_: 1.66 ± 0.30) (determined 28d post YFV exposure). Also in serum of JE-CVax- or Ixiaro^®^-vaccinated C57BL/6 mice, only nAbs against JEV (log_10_CPENT_50_: 1.66 ± 0.12 or 1. 61 ± 0.09, respectively) were detectable. Thus, neither JE-CVax nor Ixiaro^®^ induce YFV-specific nAbs in mice.

**FIG 2.**
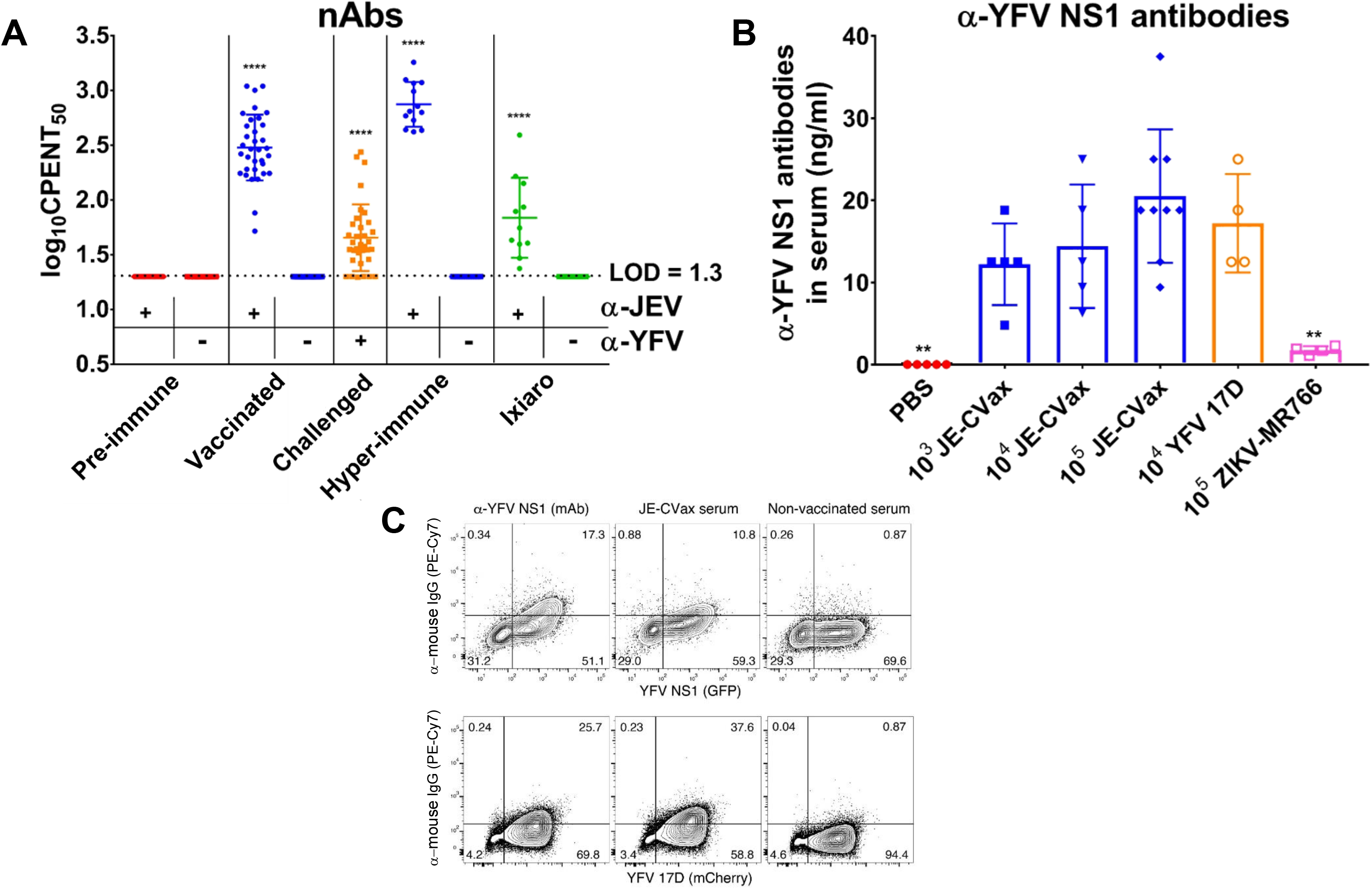
Serological analysis of serum of JE-CVax-vaccinated and YFV-17D- or ZIKV-MR766-challenged animals. **(A)** Detection of nAbs against JEV and YFV. CPE neutralization tests (CPENT) for JE-CVax (•) and YFV-17D (■) were performed on sera d0 prior to vaccination (pre-immune, red), d28 after vaccination (blue) and after challenge (study endpoint, orange) for samples of JE-CVax-vaccinated AG129 mice, of JE-CVax-vaccinated mice after subsequent YFV-17D-challenge (n = 34), of mice hyperimmunized with JE-CVax (n = 13, first bleed two weeks post last booster immunization, blue) and of mice vaccinated with Ixiaro^®^ (green). Limit of detection (LOD) for virus neutralization was log_10_20 (1.3). Data presented as log_10_CPENT_50_ (mean ± SD). The data presented are from n ≥ 3 independent experiments. Statistical significance was determined using one-way ANOVA analysis. ****p-value ≤ 0.0001 to mean log_10_CPENT_50_ titers against JEV or YFV compared to mean log_10_CPENT_50_ titers pre JE-CVax vaccination and pre YFV-17D-challenge, respectively. **(B)** Quantitation of anti-YFV NS1 binding antibodies by direct ELISA. Serum from naïve, non-vaccinated mice (red) or mice that had been vaccinated with either 10^3-5^ PFU JE-CVax (blue), or that had been infected with 10^4^ PFU YFV-17D (orange) or with 10^5^ PFU ZIKV-MR766 (pink) were collected either 28 days post immunization or when euthanized at the humane endpoint (n ≥ 5). The data shown are means of two independent analyses. Statistical significance was determined using one-way ANOVA analysis. ** p-value ≤ 0.01 compared to YFV-17D. **(C)** Binding of serum antibodies to NS1 expressing cells. HEK-293 cells were transfected with a plasmid expressing YFV-17D NS1 as a transcriptional fusion to GFP (top), or infected with the YFV-17D-mCherry reporter virus (bottom). Either 48 h after transfection or 72 h after infection, cells were stained with the anti-YFV NS1-specific mAb 1A5 (mAb, left), with serum from mice that were vaccinated with JE-CVax (center), or with serum from naïve, non-vaccinated mice (right). Graph showing flow cytometric analysis of GFP or mCherry fluorescence and visualization of anti-YFV NS1 antibody binding using a PE-Cy7 conjugated goat anti-mouse IgG secondary antibody. The fraction of NS1 positive cells (GFP or mCherry) stained by mAb 1A5 or serum of JE-CVax-immunized mice (a-mouse IgG) is given in percentage in the upper right quadrant. Data from one representative experiment out of four independent experiments.

### JE-CVax and YFV-17D induce comparable levels of anti-NS1 YFV antibodies

From the IIFA analysis (Fig. S2), it is obvious that sera from JE-CVax-vaccinated mice contain cross-reactive but non-nAbs against YFV-17D (Fig. 2A). These non-nAbs may possibly be attributed to NS1, a strong B cell antigen. To assess the presence of anti-YFV NS1 antibodies in JE-CVax-vaccinated AG129 mice, a direct ELISA was performed on sera from mice vaccinated with 10^3^-10^5^ PFU JE-CVax, or mice infected with 10^4^ PFU YFV-17D or 10^5^ PFU ZIKV-MR766. Serum was obtained either at the onset of disease (YFV-17D, 10^5^ PFU JE-CVax and ZIKV) or at 28 dpv. Levels of anti-YFV NS1 antibodies in the different JE-CVax-vaccinated groups were statistically not different (p>0.05) from the YFV-17D-infected groups (Fig. 2B). Moreover, very low cross-reactivity was noted for samples from ZIKV-infected mice. These findings were also confirmed by flow cytometry analysis (Fig. 2C). Serum antibodies from JE-CVax-vaccinated mice bound to cells that overexpress YFV NS1 as well as to cells that had been infected with YFV-17D. In fact, serum from JE-CVax-vaccinated mice resulted in a comparable staining as when a monoclonal antibody [mAb 1A5] specifically directed against YFV NS1 was used (27).

### JE-CVax induces YFV-specific antibodies mediating ADCC

To determine the potential mechanism that non-nAbs elicit for protection against YFV, an antibody-dependent cellular cytotoxicity (ADCC) reporter bioassay was carried out using YFV-17D (expressing mCherry) (28) infected HEK293T cells as target cells and murine FcγRIIIa expressing Jurkat reporter cells as effector cells (29). Hyperimmune mouse serum from JE-CVax-vaccinated AG129 mice induced clear ADCC responses and this in a dose-dependent manner, whereas serum from non-vaccinated mice failed to do so (Fig. 3A, 3B and Fig. S3A, 3B).

**FIG 3.**
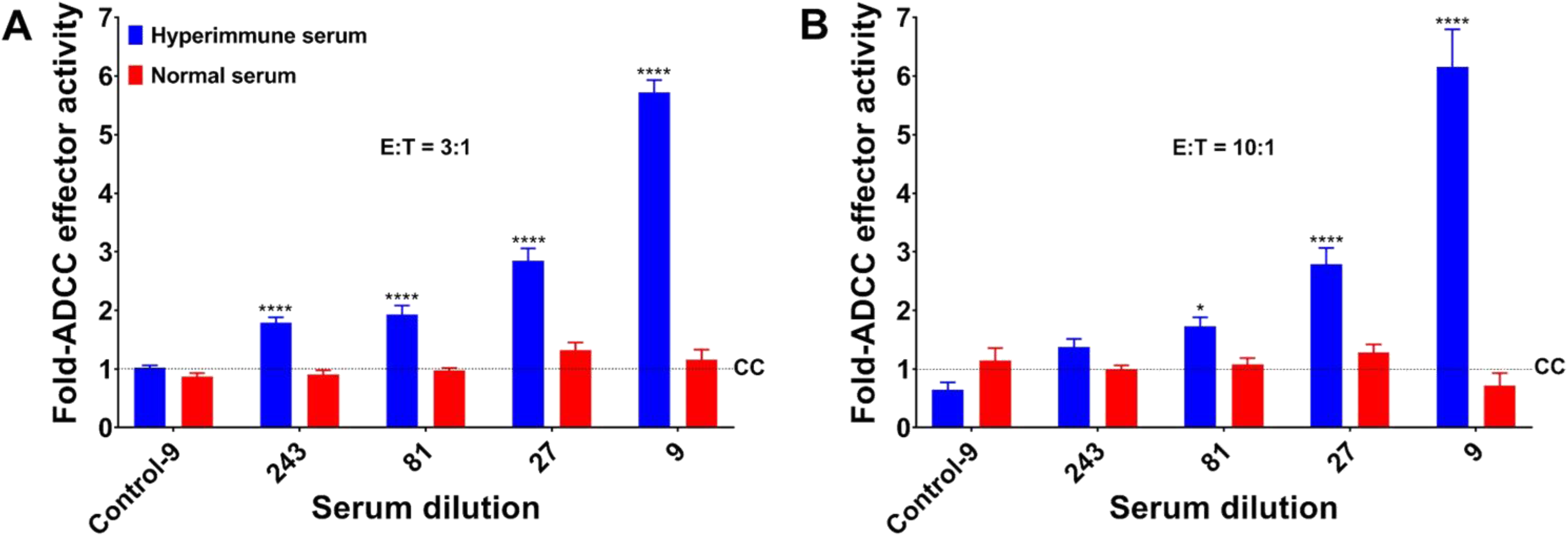
Role of antibody-dependent cell-mediated cytotoxicity (ADCC) conferred by JE-CVax hyperimmune serum in the protection against YFV. **(A, B)** JE-CVax hyperimmune serum (blue bars) was tested for its ability to mediate ADCC activity compared to serum of non-vaccinated mice (normal serum, red bars) at 3:1 **(A)** and 10:1 **(B)** effector (E): target (T) cell ratios. Experiments were conducted twice, each in triplicate, and data presented as mean ± SEM as fold changes compared to control (CC) [i.e. mean reporter signal plus three standard deviations from E:T in the absence of hyperimmune serum]. Values from non-infected target cells incubated with E in the presence of either hyperimmune serum or normal serum at highest antibody concentrations (dilution 1:9) are indicated as Control-9. Statistical significance was determined using two-way ANOVA analysis. *, **** p-value ≤ 0.05 and ≤ 0.0001 compared to normal serum.

### JE-CVax induces polyfunctional T cell responses against both YFV and JEV antigens

To assess whether also cellular immune response against YFV may contribute to the protective activity, ELISpot assays (TNF-α and/or INF-γ) and intracellular staining of cytokines (TNF-α and INF-γ) were performed on total splenocytes obtained from AG129 mice (n = 5) and C57BL/6 mice (n = 10) 18- and 4-weeks after JE-CVax immunization, respectively. Unlike splenocytes of non-vaccinated mice (Fig. S7), robust and specific IFN-γ and/or TNF-α production was observed from splenocytes of either mouse strain vaccinated with JE-CVax (Fig. 4A-C and Fig. S4) after recall with both, an MHC class I-restricted peptide derived from YFV-17D NS3 or YFV-17D total cellular antigen. In line, flow cytometric analysis revealed robust and YFV-specific intracellular cytokine production in CD4^+^ and CD8^+^ T cells from spleens of JE-CVax-vaccinated AG129 and C57BL/6 mice when stimulated ex vivo with the YFV NS3 peptide or YFV-17D total cellular antigen (Fig. 4D, Fig. S4–S6). Overall, JE-CVax vaccination induced specific long-lasting T cell responses against YFV and cellular immunity against YFV was more vigorous when compared to that elicited against JEV (Fig. 4A-C, Fig. S4–S6). This observation is consistent with YFV NS proteins serving as more immunogenic T cell antigens than the prM/E of JEV (30, 31).

**FIG 4.**
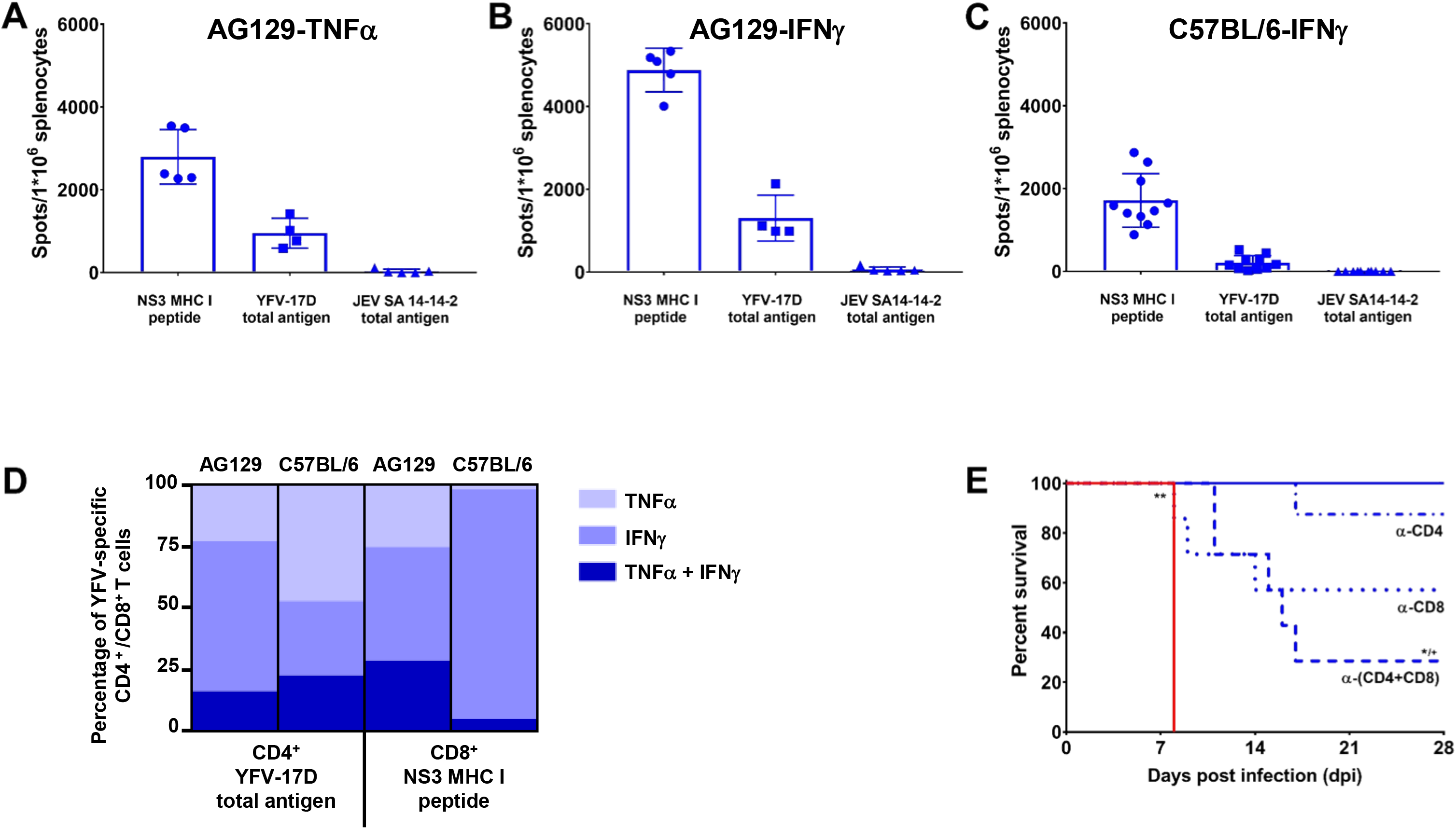
Detection of protective T cell responses directed against YFV. Detection of YFV-specific T cells. **(A-C)** ELISpot data showing TNF-α **(A)** IFN-γ **(B, C)** and production by splenocytes of AG129 mice **(A, B)** or C57BL/6 **(C)** at 18- and 4-weeks, respectively, post-vaccination with 10^4^ PFU of JE-CVax, following 16 h *ex vivo* re-stimulation with either an MHC class I restricted peptide derived from YFV-17D NS3 (32) or the lysate of YFV-17D- or JEV SA14-14-2-infected Vero E6 cells. Stimulation using lysate of non infected Vero E6 cells served as negative control. Spot counts for TNF-α **(A)** IFN-γ **(B, C)** or producing cells from **(A, B)** AG129 mice (n = 5) or (c) C57BL/6 mice (n = 10) animals, respectively. The data shown are derived from two independent experiments. Spot counts were normalized by subtraction of the number of spots in corresponding wells stimulated with uninfected Vero E6 cell lysate). **(D)** Cytokine expression profile of YFV-specific T cells. IFN-γ and TNF-α production profile of YFV-specific CD4^+^ or CD8^+^ T cells from JE-CVax-vaccinated AG129 or C57BL/6, 18- and 4-weeks, respectively, postvaccination, as determined by intracellular cytokine staining. Mouse splenocytes were stimulated 16h *ex vivo* with either an MHC I restricted NS3 peptide, cell lysate of YFV-17D-infected Vero E6 cells or uninfected Vero E6 cells. The data shown are derived from two independent experiments and normalized by subtraction of number of cytokine-secreting T cells in corresponding samples in which uninfected Vero E6 cell lysate was used as recall antigen. **(E)** T cell-mediated *in vivo* protection against YFV. Loss of protection resulting from antibody-mediated T cell depletion (30, 52) suggests a direct functional involvement of CD4^+^ and CD8^+^ T cells in JE-CVax-mediated immunity against YFV in C57BL/6 mice (n≥7) that had been vaccinated with 10^4^ PFU of JE-CVax and subsequently challenged intracranially with 10^4^ PFU of YFV-17D. Depletions were performed by administration of 0.5 mg of a-mouse CD4 and/or a-mouse CD8a antibodies i.p. on day (−2) and day 0 each prior to YFV challenge. Log-rank (Mantel-Cox) survival analysis test was performed for statistical significance. *, ** p-value ≤ 0.05 and ≤ 0.01 compared to vaccinated group (n=5) and ^+^ p-value ≤ 0.05 compared to CD4^+^ depleted group (n=8).

### Both CD4^+^ and CD8^+^ T cells contribute to JE-CVax-mediated protection against YFV

To determine whether YFV-specific T cell responses directly contribute to the JE-CVax-mediated protection against YFV, T cell depletion experiments were performed in C57BL/6 mice (32). Five-week post-vaccination animals were administered with anti-mouse CD4 and/or anti-mouse CD8a T cell-depleting antibodies twice, i.e. two days before and immediately prior to YFV challenge. Unlike for vaccinated (but not further treated) animals [that were included as immunization controls (n=5) and that survived an intracranial YFV challenge], in vaccinated but antibody-treated mice the previously observed full protection against YFV was partially lost by targeting CD4^+^ (n=1/8), CD8^+^ (n=3/7) and CD4^++^CD8^+^ (n=5/7) T cells for depletion (Fig. 4E). All non-vaccinated animals (n=5, p-value; 0.0027) succumbed to YFV challenge as before (Fig. 1D). The mortality resulting from T cell depletion, especially the increased mortality observed in the double-depleted animal group, suggests that in the absence of nAb both CD4^+^ and CD8^+^ T cells contribute to the JE-CVax-mediated protection against YFV in C57BL/6 mice.

## DISCUSSION

Neutralizing antibodies against the E protein are generally considered as a primary correlate of protection against flaviviruses (5, 15). However, some preclinical studies suggested that several NS proteins when used as immunogen for vaccination alone or in combination could induce some degree of protection in mice and non-human primates against viral challenge (17–20). For instance, a recombinant vaccinia virus or replication deficient adenoviral vectors expressing YFV-17D NS1, NS2a and NS2b together or NS3 alone, respectively, resulted in some partial protective immunity against a lethal challenge of YFV-17D in mice (17, 23). However, full survival could never be achieved and reached maximally 60-80% versus 100% for YFV-17D vaccinated controls. Similarly, 80% protection against challenge with the African YFV strain Dakar 1279 was observed in monkeys following repeated immunization with purified NS1 as sole vaccine antigen and protection correlated with the levels of non-nAbs against NS1 (19). Conversely, vaccination of mice with E alone (or in combination with NS1 or NS3) resulted in complete protection (23, 24).

The live-attenuated JE-CVax vaccine is a chimeric flavivirus that consist of the genome of YFV-17D from which the prM/E genes have been replaced by those of JEV SA14-14-2. We, therefore, used JE-CVax to assess whether it can offer, besides protection against JEV, also protection against YFV challenge. Since AG129 mice are highly susceptible to lethal JEV SA14-14-2 and YFV-17D infection (Fig. S8, (28, 33)), we used these two vaccine strains as established surrogates for wt-JEV (33) and wt-YFV (34, 35), respectively (demanding lower biosafety containment for handling). A single dose of JE-CVax provided in addition to the expected protection against JEV challenge, a near to complete protection (35/36) against a massive 1000x-LD_50_ challenge with YFV (Fig. 1B). The protective activity against YFV was raised fast, and a survival benefit could be observed already within 7 days after vaccination (Fig. 1C). These observations were further corroborated by the fact that vaccination of immunocompetent C57BL/6 mice with a single dose of JE-CVax provided complete protection (21/21) against lethal intracranial challenge (17, 23) with YFV (Fig. 1D and Fig 4E). This activity of JE-CVax was YFV-specific, as AG129 mice that had been vaccinated with Ixiaro^®^ (the inactivated JEV vaccine) were not protected against lethal YFV challenge. Furthermore, mice that had been vaccinated with JE-CVax and that later survived YFV challenge did not survive a subsequent lethal challenge with the ZIKV (Fig. S1).

Others have shown in mice that cross-reactive antibodies (together with cross-reactive T cells) may confer partial protection against flaviviruses from different serocomplexes as demonstrated for JEV (both Vero cell-derived JEV-P3 strain based inactivated vaccine and JEV SA14-14-2) vaccinated mice challenged with DENV (26). Likewise, a chimeric Japanese encephalitis/dengue-2 virus experimental vaccine ChinDENV (originally designed to induce immunity against DENV-2 prM/E) was shown to protect against both JEV and DENV-2 challenge in mice (31, 36). The protection observed could however still largely be explained by the induction of considerable levels of E-specific partially cross-reactive nAbs that neutralized both DENV and JEV. Additionally, the study (31) demonstrated that vaccination also induces JEV antigen-specific T cell responses (T cells producing IFN-γ and IL-2 following stimulation with JEV antigen), suggesting a possible contribution of the cellular immunity in the defence against JEV challenge in ChinDENV-immunized mice. Another study, in which AG129 and IFNAR mice were used, reported that during heterotypic dengue virus infection, CD8^+^ T-cells provide some degree of protection in the absence of detectable levels of nAbs (37). However, in this case a mild (sublethal) DENV-4 infection was used for priming rather than a true vaccine and despite this priming, only limited (partial) and short-term protection was observed against DENV-2. Moreover, the E-proteins of DENV-2 and DENV-4 share high (∼64%) sequence homology, including conserved T cell epitopes (38) and therefore, the contribution of the E-protein for protective T cell responses could obviously not be distinguished from what DENV NS proteins may contribute. By contrast, although we also observed some residual cross-reactivity for YFV in serum samples of JE-CVax-vaccinated mice (binding antibody in IIFA, Fig. S2), JE-CVax failed to induce any detectable nAb titers against YFV even following repeated boosting (Fig. 2A). This finding is in full accordance with the absence of cross-nAbs in mice, monkeys and humans post JE-CVax or YFV-17D vaccinations (39–41). Therefore, cross-reactivity of serum from JE-CVax-vaccinated mice could be attributed to (a) induction of E-based broad flavivirus cross-reactive non-nAbs resulting from flavivirus infection/vaccination (26, 42) and (b) reactivity against YFV-NS1 that is expressed within and on the surface of infected cells and is target of specific binding but non-nAbs (20, 43). In fact, we demonstrate equivalent levels of anti-YFV-NS1 antibodies in either JE-CVax-vaccinated or YFV-17D-infected mice (Fig. 2B-C).

Although, we show that JE-CVax immunization resulted in complete protection against YFV-induced disease, there were variations in the actual levels of anti-YFV nAbs post YFV challenge. In some animals, no nAbs were detected against YFV post YFV-17D-challenge. This lack of YFV nAb provides strong evidence that JE-CVax even conferred sterilizing immunity in these mice. Since, such protection cannot be explained by nAb against the YFV-prM/E, it may be accredited to non-nAbs (42–44) and/or adaptive cellular immunity (16, 30, 32, 45). Some correlation between anti-NS1 antibody levels and the dose of JE-CVax needed to cause protection was observed (Fig. 2B.C and Fig. S1A) and serum from JE-CVax-vaccinated mice was found to induce an ADCC response against YFV-17D (Fig. 3 and Fig. S3). In addition, in our model both CD4^+^ and CD8^+^ T cells seem (in a likely association with anti-NS1 antibodies) to be involved in JE-CVax-mediated protection against YFV (Fig. 4E). Previously, only humoral immunity and CD4^+^ T (but not CD8^+^ T) cells have been implied to be sufficient and required for protection against YFV (16), with a strong emphasis on nAb as historically established immunological correlate of protection for YFV (5) and for flaviviruses (such as JEV) in general (1, 15, 39). NAbs possibly block viral spread, whereas cellular immunity efficiently eliminates intracellular viruses either directly, or targeted by non-nAbs towards infected cells in an Fc-dependent manner (for example, via ADCC towards cells expressing YFV NS1 on their surface) (18, 21, 43, 44). Indeed, YF-induced CD8^+^ T cells have been shown to act as a ‘backup defence’ system in the absence of nAbs and to participate in viral clearance in particular from the CNS in mice (23, 32). Moreover, strong CD8^+^ T cell responses are also detected in the response to human vaccines (30). As we show here, immunization of AG129 and C57BL/6 mice with JE-CVax elicited protective polyfunctional YFV antigen-specific CD4^+^ and CD8^+^ T cell responses (expressing the Th1-type cytokines TNF-α and IFN-γ), which is in line with a previous study with a chimeric Japanese encephalitis/dengue-2 virus vaccine (31). Collectively, our data suggest that JE-CVax mediated vigorous protection against lethal YFV challenge depends on the combined effect of several effector principles, including both the humoral and cellular immune responses, yet definitely other than nAbs.

To assess the efficacy of JE-CVax, we employed mice as *in vivo* model, and the YFV-17D vaccine strain (16, 17, 23, 44, 46) as challenge virus. This experimental setup implies some constraints. Generally, mice are not susceptible to human flavivirus infection (47) and the YFV-17D is not virulent in humans (5). To overcome some of these limitation, we made use of two established, complementary and stringent mouse infection models that are accepted surrogates for testing of flavivirus vaccines (11); (i) immunocompromised mice (AG129 mice; (28, 34, 35)) that develop fatal neurotropic infection when challenged peripherally with YFV-17D, in particular when inoculated with a highly lethal (1000x-LD_50_) challenge virus dose (Fig. S8, (28)), and (ii) immunocompetent wt mice (C57BL/6; (16, 17, 23)) that can be challenged i.c. with YFV-17D. YFV-17D was originally developed by adapting a viscerotropic clinical isolate (YFV-Asibi) to replication in mouse brains (‘fixed to mouse brain’) (48). This vaccine virus can hence, be considered as a genuine mouse adapted neurotropic and neurovirulent YFV strain. For this reason, it could also be considered in this study as the challenge virus. Besides experimental convenience (YFV-17D does not require BSL3 containment), YFV-17D is a widely accepted, i.e. best characterized and hence most widely used challenge strain in mouse models. Classical i.c. inoculation of YFV-17D consistently causes fatal disease in mice (46) that cannot be distinguished from that induced by wt-YFV strains (44, 46). Also, because JE-CVax expresses the prM/E of JEV that belongs to another serocomplex than YFV, vaccination and subsequent challenge with YFV-17D compares to a certain extent to a heterotypic flavivirus vaccination-challenge (as described by others (37)), however with a markedly more pronounced vaccine efficacy. Therefore, in conclusion, similar mechanisms should hold when using wild-type YFV as the challenge virus (44, 46). Obviously, before proceeding to clinical use, this principle should be confirmed in JE-CVax vaccinated non-human primates demonstrating protection from subsequent challenge with virulent wt-YFV strains.

Previous safety and immunogenicity studies of chimeric live-attenuated viruses (JE-CVax and ChimeriVax-DEN2) in humans indicated that pre-existing immunity to the parental YFV-17D vaccine virus does not interfere with immunization, but rather induces long-lasting cross-neutralizing antibody responses (39, 49, 50). Importantly, if our data on the dual protection conferred by c-LAV in mice can be translated to other species, including humans, this implies that the JE vaccine JE-CVax (and likewise Imojev^®^) may provide dual protection, i.e. against both JEV and YFV. A dual protective effect may be of particular relevance in case YFV may one day – as is feared (8, 9) – cause outbreaks in (sub)tropical Asia. Given the capacity problems with the production of the current YFV vaccine, having another licensed vaccine (i.e. JE-CVax/Imojev^®^) available as an alternative means to protect against YFV may at such time help to contain an outbreak. In addition, those already vaccinated with JE-CVax may be expected to be protected against YFV. A similar principle may apply to other chimeric flavivirus vaccines that consist of a YFV-17D backbone (such as CYD-TDV/Dengvaxia^®^) (12) and others under development (such as our recently proposed chimeric YFV-17D/ZIKV vaccine candidate) (28). Likewise, c-LAVs could be generated against DENV and other viruses that may cause using the backbone of the parent virus (e.g. DENV) from which the prM/E genes have been replaced by that of antigenically more distantly related viruses or serotypes. Such approach may avoid potentially harmful nAb responses. The same principle may apply to c-LAV for ZIKV using a ZIKV backbone (51) and prM/E sequences of another flavivirus that is not shown to cause ADE.

To conclude, we demonstrate that JE-CVax efficiently and rapidly induces high cross-protective efficacy (∼100%) in mice against YFV challenge, even with an exceedingly aggressive challenge inoculum. The study provides evidence that c-LAV flavivirus vaccines may be developed solely based on NS proteins. Moreover, immunization with a chimeric flavivirus, whereby the prM/E genes of the backbone have been replaced by that of yet another flavivirus may have a dual protective effect. A vaccine, such as Imojev^®^/JE-CVax may thus be suitable for off-label use, namely for protection against YFV, which in this case is not mediated by nAbs.

## MATERIALS and METHODS

### Cells and medium

BHK-21J and Vero E6 cells used in this study were a generous gift from Peter Bredenbeek, LUMC, NL. Cells were maintained in seeding medium containing MEM Rega-3 medium (Gibco, Belgium) supplemented with 10% fetal calf serum (FCS, Gibco, Belgium), 2mM glutamine (Gibco, Belgium) and 0.75% sodium bicarbonate (Gibco, Belgium). HEK 293 (human embryonic kidney 293 cells; ATCC CRL-1573) cells were cultured in DMEM (Gibco, Belgium), containing 10% FCS and 100 units/ml penicillin-streptomycin solution (Pen/Strep, Gibco, Belgium). Virus culture and cytopathic effect-based virus neutralization assays (CPENT) were performed in assay medium, which is the seeding medium supplemented with only 2% FCS. HEK 293 cells were transfected with YF-NS1-GFP using TransIT^®^-LT1 transfection reagent (Mirus Bio LLC, Belgium), according to the manufacturer’s instructions. Infection of HEK 293 cells with YFV-17D-mCherry (see below) was performed in DMEM medium supplemented with 2% FCS and 100 units/ml Pen/Strep solution. All cultures were maintained at 37°C in an atmosphere of 5% CO2 and 95%–99% humidity.

### Virus, vaccines and antigens

Stamaril^®^ (G5400) and Ixiaro^®^ (JEV16F290) were from Sanofi Pasteur (France) and Valneva (Austria), respectively. Stamaril^®^ was passaged two times in Vero E6 cells (YFV-17D-G5400P2), and stored at (−80°C). YFV-17D-G5400P2 was used throughout the study to challenge mice and is referred to as YFV-17D. YFV-17D-based Japanese encephalitis c-LAV Imojev^®^ (Chimerivax-JE, JE-CVax) is not available in Europe and was, therefore, retro-engineered according to publicly available sequence information (52) (Patent Number: WO2006044857A2). To that end, a DNA fragment encoding the prM and E proteins of JEV vaccine strain SA14-14-2 was custom synthetized (IDT Integrated DNA Technologies, Haasrode, Belgium) and subcloned into the YFV-17D expression vector pShuttle-YFV-17D (52) (Patent Number: WO2014174078 A1) of our Plasmid-Launched Live-Attenuated Vaccine (PLLAV)-YFV-17D platform using standard molecular biology techniques and thereby replacing the YFV-17D prM/E sequences. Two adaptive mutations in NS2A and NS4B genes and an additional Kas1 site at the end of the prM/E coding sequence (52) were introduced by site-directed mutagenesis. To generate JE-CVax virus, BHK 21J cells were transfected with PLLAV-JE-CVax using TransIT^®^-LT1 transfection reagent, following the manufacturer’s instructions. Upon onset of CPE, JE-CVax virus was harvested, centrifuged at 4000 rpm at 4°C for 10 minutes, aliquoted and stored at (−80°C). Similarly, the live-attenuated Japanese encephalitis virus vaccine JEV SA14-14-2 (Genbank AF315119.1) was generated fully synthetically from overlapping DNA fragments (IDT Integrated DNA Technologies, Haasrode, Belgium), assembled by overlap-extension PCR and subsequent homologous recombination in yeast. The recombinant JE-CVax and JEV SA14-14-2 viruses were subsequently passaged on Vero E6 cells to generate virus stocks. As an alternative challenge virus, ZIKV strain MR766 was used (53). Virus titers were determined on BHK-21J cells by plaque assays (plaque forming units/ml; PFU/ml) and CPE-based assays (TCID_50_/ml) as described below.

A YFV-17D reporter virus (YFV-17D-mCherry) was generated that expresses the red-fluorescent protein mCherry as a translational fusion to the N-terminus of YFV-17D C protein. In brief, using standard PCR techniques and homologous recombination in yeast a synthetic DNA fragment encoding codons 2-236 of mCherry (Genbank AY678264.1) was inserted in YFV-17D genome (54) immediately downstream of nucleotide position 181, flanked (i) at its 5’ terminus by a glycine-serine linker (BamH1 site), and (ii) at the 3’ end by a *Thosea asigna* virus 2A peptide (EGRGSLLTCGDVEENPG/P) (55) followed by a repeat of codons 2-21 of YFV-17D C gene, yet with an alternative codon usage to avoid RNA recombination during viral replication. YFV-17D-mCherry was rescued by transfection of the resulting PLLAV-YFV-17D-mCherry of BHK 21J as before (28). A full characterization of YFV-17D-mCherry is given elsewhere (Schmid et al., in preparation).

Plasmid pCMV-YFV-17D NS1-IRES-EGFP that drives the mammalian expression of YFV-17D NS1 as a transcriptional fusion to EGFP was generated by PCR cloning of YFV-17D nt 2381–3508 cDNA (including an E protein derived N-terminal signal peptide) (56) flanked by a 5’ terminal Kozak sequence and 3’ terminal stop codon into the Nhe1 and Sal1 sites of pIRES2-EGFP (Clontech cat # 6029-1).

An MHC I class-restricted peptide from YFV-17D non-structural protein 3 (NS3) (sequence ATLTYRML) (57) was synthetized by Eurogentec (Seraing, Belgium). Total cellular antigen for YFV-17D and JEV SA14-14-2 was prepared first by infecting Vero E6 cells with 0.1 MOI YFV-17D or JEV SA14-14-2, respectively. Non-infected Vero E6 cells were used as control. Four days post-infection, cells were harvested either by trypsinization or by detaching through pipetting, when a cytopathic effect (CPE) was visible. Following centrifugation, cell pellets were resuspended in PBS and cell lysates were prepared by four freeze-thaw cycles. Overnight UV-irradiation was performed to inactivate the virus in the cell lysate preparations and large debris was removed via filtering through 70 μm cell strainers (BD Biosciences).

### Animals, hyper-immune serum, infection and T cell depletions

Host IFN signalling is the major barrier to the viscerotropism and for pathogenicity of neurotropic flaviviruses (58). In line, wild-type mice are poorly susceptible to infection with flaviviruses (28, 59–61) including YFV-17D infection/vaccination (34, 35, 62). Likewise, type I (IFN-α/β) and type II (IFN-γ) interferon signalling mutually controls YFV-17D infection. Unlike humans (63, 64), type I IFN (IFN-α/β) can restrict YFV-Asibi as well as YFV-17D infection in mice (34, 35, 62, 65). Similarly, IFN-γ exerts restriction on YFV-17D replication, dissemination and clearance in mice (35, 66). YFV-17D is neurotropic in wt-mice when directly injected into the brain (46). In AG129 mice, INF-α/β and γ receptors are knocked out, which results in neurotropic infection following peripheral inoculation of YFV-17D. Therefore, to allow transient replication (and thus vaccination) of YFV-17D in wt-mice, MAR1-5A3 antibodies were co-administered to block type-1 IFN signalling into C57BL/6 mice.

Immunodeficient interferon (IFN)-α/β and -γ receptor knockout mice (AG129; B&K Universal, Marshall Bio resources, UK) were bred in-house. AG129 mice have been shown to be highly susceptible to lethal YFV-17D infection serving as a well-established surrogate rodent challenge model for wt-YFV infection (34, 35, 67). Six to eight weeks old male AG129 mice were used for all experiments after random assignment into different groups. Animals were kept in individually ventilated type-2 filter top cages on a 12 hour day/night cycle with water and food *ad libitum.* Housing of animals and procedures involving animal experimentation were conducted in accordance with institutional guidelines approved by the Ethical Committee of the KU Leuven, Belgium [licenses P168/2012, P103/2015, P140/2016 and P005/2018]. Throughout the study, animals were vaccinated intraperitoneally (i.p.) with either 10^4^ plaque forming units (PFU) of JE-CVax, 1/6^th^ human doses of Ixiaro^®^ (25) or 2% assay medium. Animals vaccinated with Ixiaro^®^ were boosted on 14 dpv with 1/6^th^ human dose of Ixiaro^®^. All the vaccinated animals were challenged i.p. with either 10^3^ PFU of YFV-17D or JEV SA14-14-2 (both corresponding to 1000 LD_50_) 28 dpv, if not stated otherwise. An additional four weeks post YFV-17D challenge, some animals were re-challenged i.p. with 10^4^ PFU ZIKV-MR776 (53). Animals were observed for morbidity (weight loss) and humane endpoint once daily. The humane endpoint is defined as either paresis/difficulty in walking, paralysis (hind legs/soured eyes), moribundity / ataxia / tremors / difficulty in breathing, 20% weight loss or quick weight loss (15% within 1 or 2 days) and animals were immediately euthanized once a humane endpoint was reached. Throughout the study, bleedings were performed through submandibular puncture on day 0 (pre-vaccinated), day 28 (post-vaccinated) and day 56 (post-challenged).

Hyper-immune serum was prepared by vaccinating AG129 mice with 10^4^ PFU JE-CVax, followed by two boosts with 10^5^ PFU JE-CVax in 14-day intervals. Another 14 days after the second booster, animals were bled twice per week for the following four weeks. All serum batches were then pooled and CPENT assays for JE-CVax and YFV-17D were performed. We did not observe any YFV nAbs in the hyperimmune-sera but did see a ∼3.6-fold (log_10_CPENT_50_ titers 3.03 ± 0.18) selective increase in neutralizing titers against JE-CVax compared to single vaccination. (Fig. 3A). Normal mouse control serum was prepared by pooling serum from 18 non-vaccinated AG129 mice.

Immunocompetent wt-C57BL/6JOIaHsd i.e. C57BL/6 were purchased from ENVIGO Labs, Netherlands and were maintained and manipulated as described for AG129 mice with some modifications (23, 28). Since, flaviviruses do not readily replicate in immunocompetent wild type mice (28), they were immunized with 10^4^ PFU JE-CVax in the presence of 2.5 mg of an IFN alpha/beta receptor subunit 1 (IFNAR-1) binding monoclonal antibody, MAR1-5A3, administered i.p. one day prior to immunization. 0.5 mg MAR1-5A3 antibody was also re-administered i.p. on day 4 and day 7 post-vaccination. Animals were bled 28 days post-vaccination and challenged i.c. with 10^4^ PFU YFV-17D in 30 μL of volume. A full characterization of immunogenicity of YFV-17D in various mouse strains is given elsewhere (Ma et al., in preparation). For T cell depletion studies, C57BL/6 mice were either sham-vaccinated or vaccinated i.p. with 1×10^4^ PFU of JE-CVax 35 days prior to i.c. challenge with 1×10^4^ PFU of YFV-17D. At day (−2) and day 0 prior to YFV challenge, 0.5 mg of either anti-mouse CD4 (Clone GK1.5, Leinco Technologies, USA) or anti-mouse CD8a (Clone 53-6.7, Leinco Technologies, USA) or a combination of both was administered i.p. (32, 68).

### Indirect immunofluorescence assay (IIFA)

To determine the seroconversion of animals, all JEV, YFV, and ZIKV IgG-IIFAs were performed as per the manufacturer’s instruction (Euroimmune, Lübeck, Germany), except for the use of labelled secondary antibody and the mounting agent glycerine, which were replaced by Alexa Fluor 488 goat anti-mouse IgG (A-11029, ThermoFisher Scientific) and DAPI (ProLong^®^ Antifade Reagent with DAPI, ThermoFisher Scientific), respectively. Serum from nonvaccinated animals served as naïve, negative control. Slides were visualized using a fluorescence microscope (FLoid Cell Imaging Station, ThermoFisher Scientific).

### Plaque assay and plaque reduction neutralization test (PRNT)

Viral titers of YFV-17D or JE-CVax preparations were determined using plaque assays on BHK-21J cells. In brief, 10^6^ BHK-21J cells per well were plated in 6-well plates and cultured overnight in seeding medium. Cells were washed with PBS and inoculated with virus of different dilutions prepared in the assay medium for one hour at room temperature (RT). Culture supernatants of uninfected cells were used as negative controls. Cells were thoroughly washed with the assay medium and overlaid with MEM-2X (Gibco, Belgium) supplemented with 4% FCS and 0.75% sodium bicarbonate containing 0.5% low melting agarose (Invitrogen, USA). The overlay was allowed to solidify at RT, cells were then cultured for 7 days at 37 °C, fixed with 8% formaldehyde and stained with methylene blue. Plaques were manually counted and plaque titer was determined as PFU/ml.

Throughout the study, all the virus neutralization assays i.e. PRNT and CPE-based virus neutralization assay (CPENT) were performed with YFV-17D and JE-CVax. JE-CVax has previously been established as a safe substitute for JEV, a BSL-3 pathogen, when virus neutralization tests need to be performed in BSL-2 (69). PRNT assays were performed in a similar way as the plaque assays for viral titration with some modifications. Briefly, an additional step was added, where different serum dilutions made in the assay medium were first inoculated with YFV-17D (20-50 PFU) or JE-CVax (50-100 PFU) virus for 1 h at 37 °C and then added to the cells. All sera were assayed in triplicate in serial dilutions 1:20, 1;66, 1:200, 1:660, 1:2000 and 1:6600. Plaques were manually counted and PRNT_50_ were calculated using the Reed and Muench method (70). Culture-derived YFV-17D or JE-CVax were used as positive virus controls, while culture supernatants of uninfected cells were used as negative cell control. PRNT_50_ values for each sample represent geometric means of three independent repeats and data presented as (mean ^+^ SD) log_10_PRNT_50_.

### Cell-based cytopathic (CPE) assay and CPE-based virus neutralization test (CPENT)

Viral titers for culture-derived YFV-17D or JE-CVax (TCID_50_) and 50% neutralizing antibody titers (log_10_CPENT_50_) were determined using cytopathic effect (CPE)-based cell assays and CPE-based virus neutralization tests (CPENT), respectively, on BHK-21J cells (71) with some modifications. In brief, 2 × 10^4^ BHK-21J cells/well were plated in 96-well plates overnight in seeding medium. The medium was then replaced with assay medium containing different virus dilutions and cultured for 5 days at 37°C. Later, assays were first visually scored for CPE and then stained with MTS/Phenazine methosulphate (PMS; Sigma-Aldrich) solution for 1.5 h at 37 °C in the dark. Post MTS/PMS staining absorbance was measured at 498 nm for each well. All assays were performed in six replicates and TCID_50_/ml was determined using the Reed and Muench method (70).

CPENT assays were performed in a similar way as the CPE assays for viral titration with some modifications. Briefly, an additional step was added, where different serum dilutions made in the assay medium were first inoculated with 100 TCID_50_ YFV-17D or JE-CVax virus for 1 h at 37 °C and then added to the cells. All sera were assayed in triplicate in serial dilutions 1:20, 1;66, 1:200, 1:660, 1:2000 and 1:6600. CPE neutralization was calculated with the following formula: % neutralization activity = % CPE reduction = (OD_Virus+Serum_ – OD_vc_) *100 / (OD_cc_ – OD_vc_) and 50% neutralization titers (CPENT_50_) were calculated using the Reed and Muench method (70). Culture-derived YFV-17D or JE-CVax were used as positive virus controls, while culture supernatants of uninfected cells were used as negative cell control. CPENT_50_ values for each sample represent geometric means of three independent repeats and data presented as (mean ± SD) log_10_CPENT_50_. The CPENT assay for detection of nAbs was validated against a standard PRNT, yielding a strong correlation (R^2^ = 0.71; p = 0.018) between PRNT_50_ and CPENT_50_ (Fig. S9A) and similar levels of anti-JEV nAbs titers in post-vaccination and post-challenge serum samples (Fig. S9B, 9C).

### Enzyme-linked immunosorbent assay (ELISA)

Serum antibodies recognizing YFV NS1 were detected by indirect ELISA, in essence, as previously described (72, 73). In brief, ELISA plates (Nunc MaxiSorp, ThermoFisher Scientific) were coated with 1 μg/ml recombinant YFV NS1 (Biorad, cat # PIP052A) in 50 mM carbonate buffer (pH; 9.6) overnight at 4°C. After three washes with PBS-T (PBS with 0,05% Tween 80), plates were blocked with 2% BSA in PBS-T for 1 h at 37°C, or alternatively overnight at 4°C. After three washes with PBS-T, wells were treated with serial dilutions of test sera (2-fold serial dilution in PBS-T) for 2 h at room temperature. Serial dilutions of the YFV NS1-specific mouse IgG2a monoclonal antibody (clone 1A5, kindly provided by J.J. Schlesinger) (27) starting at 10 μg/mL served as standard. After four washes with PBS, plates were incubated with horseradish peroxidase-labelled goat anti-mouse IgG antibody (Sigma-Aldrich, cat # AP124P, diluted 1:3000 in PBS-T) for 1 h. After another four washes with PBS, bound antibodies were detected via conversion of added TMB (SureBlue TMB Microwell Peroxidase; KPL). The reaction was stopped after 10 minutes by adding equal quantities of 1 M HCl solution, and absorbance was measured at 450 nm. After background subtraction, relative anti-YFV NS1 titers were determined by comparison to the standard curve generated for mAb 1A5 included in each assay plate. To that end, the dilution at which each individual test serum yielded an OD 450 of 1 was used to calculate an absolute anti-NS1 antibody concentration (equivalent concentration), assuming a similar binding to YFV-17D NS1 as by mAb 1A5. Only values that exceeded three times the background signal were considered positive.

### Antibody-dependent cell-mediated cytotoxicity (ADCC) bioassay

To assess the possible role of non-neutralizing YFV antibody-mediated protection against YFV post JE-CVax vaccination, antibody-dependent cell-mediated cytotoxicity (ADCC) bioassays (29) were performed as prescribed by the manufacturer (ADCC reporter bioassay, complete kit, Promega, cat # G7010). In brief, target cells (T) were prepared by infecting HEK 293T cells with YFV-17D-mCherry in assay medium. Cells were incubated at 37°C post-infection and later upon onset of CPE harvested by trypsinization. Cells were again plated in white, flat-bottom 96-well assay plates (Viewplate-96, PerkinElmer cat # 6005181) at a density of 7500 and 25000 cells per well for 8 hrs at 37°C in assay medium. Later, in a separate 96 well plate, JE-CVax hyper-immune and non-immune heat inactivated mouse serum samples (starting dilution of 1/9) were serially diluted 3-folds in RPMI 1640 medium (Gibco, ThermoFisher Scientific) supplemented with 4% low IgG serum. ADCC bioassay effector cells (Jurkat V variant cells) were diluted to 3 × 10^6^ cells/ml in RPMI 1640 medium. The supernatant from the infected cell plate was replaced with fresh RPMI medium (25 μl/well) and diluted serum samples (25 μl/well) and E cells (25 μl/well) were added to the infection plate. After incubation at 37°C for 24 h, Bio-Glo luciferase assay reagent (75 μl/well) was added, and luminescence was measured using a Spark^®^ Multimode Microplate Reader (Tecan). The average background plus three standard deviations was calculated and used as background.

### Intracellular staining of NS1 protein in HEK-293 cells

HEK-293 cells transfected with pCMV-YFV-17D NS1-IRES-EGFP or infected with YFV-17D-mCherry were detached with trypsin-EDTA (0,05%), centrifuged (at 2500 rpm and 4°C for 5 minutes) and suspended in FACS-B (DPBS, no Ca^2+^/Mg^2+^, 2% FBS, 2 mM EDTA). Not more than 5 × 10^6^ cells per well were seeded into round-bottom 96-well plates (Costar, Corning Inc., Corning), spun down, and the supernatant was removed. Dead cells were stained *in vitro* with ZombieAqua (Biolegend, 1:500 diluted in DPBS) to exclude from further analysis. After washing and fixation with 2% paraformaldehyde (in FACS-B), cells were permeabilized by 0.1% saponin (in FACS-B with 1% normal mouse serum) with streptavidin added (Streptavidin/Biotin Blocking Kit, Vector Laboratories) to block endogenous biotin (permeabilizing and blocking solution). The cells were then stained with anti-NS1 primary antibody solution (clone 1A5, 5μg/ml), or JE-CVax-vaccinated mouse serum (1:10) in permeabilizing and blocking solution. Anti-NS1 antibody binding was detected by a biotinylated goat anti-mouse IgG secondary antibody solution (ThermoFisher Scientific, cat # A16076; 1:200 dilution). The biotinylated secondary antibody was stained subsequently with streptavidin-PE-Cy7 (Biolegend, 1:200 dilution). After washing the cells, they were resuspended in FACS-B and filtered through 100 μm nylon meshes (Sefar, ELKO Filtering, 03-100/44) prior to analysis on a flow cytometer (LSR Fortessa X-20, Becton Dickinson). The data were analysed using FlowJo 10 software (TreeStar). The gating strategy for the analysis is depicted in Fig. S10A.

### Processing of mouse spleens for the preparation of single cell suspensions

Six-eight weeks old C57BL/6 or AG129 mice were vaccinated with 10^4^ PFU JE-CVax and four/eighteen weeks later, the animals were euthanized for analysis. Spleens were harvested and processed for ELISpot and flow cytometric analysis. To generate single-cell suspensions, spleens were pushed through 70 μm cell strainers (BD Biosciences) with syringe plungers, digested in 1.0 mg/ml type-1 collagenase and 10 U/ml DNase for 30 minutes, vigorously pipetted and filtered through 100 μm nylon meshes. Spleen samples were then incubated with red blood cell lysis buffer (eBioscience) for 8 minutes at room temperature and washed twice with FACS-B.

### ELISpot

Mouse TNF-α and IFN-γ Enzyme-Linked ImmunoSpot (ELISpot) assays were performed with a mouse TNF-α ELISpot kit (ImmunoSpot MTNFA-1M/5, CTL Europe GmbH) or a mouse IFN-γ ELISpot kit (ImmunoSpot MIFNG-1 M/5, CTL Europe GmbH) according to the manufacturer’s instructions. Assay plates (96-well PVDF membrane), antibodies, enzymes, substrate and diluent were included in the kits. Briefly, 4 × 10^5^ mouse splenocytes/well were plated with either with 5 μg/ml YFV-17D NS3 ATLTYRML peptide antigen (57) or with 50 μg/ml of total Vero E6 cellular antigen in RPMI 1640 medium (Gibco, Belgium) supplemented with 10% fetal bovine serum, 2mM L-glutamine and 0.75% sodium bicarbonate. After 24 hours of incubation at 37°C, spots of mouse TNF-α or IFN-γ were visualized by subsequent addition of detection antibody, enzyme and substrate. All plates were scanned using an ImmunoSpot S6 Universal Reader (CTL Europe GmbH). Spot counts were normalized by subtracting the number of spots from corresponding samples stimulated with non-infected Vero E6.

### Intracellular cytokine staining for memory T cells and flow cytometry

To restimulate memory T cells, freshly isolated single cell suspensions of splenocytes were seeded at 3 × 10^6^ cells density per well in a round-bottom 96-well plate, and incubated with either 5 μg/ml of the MHC I class restricted peptide from YFV-17D NS3 (ATLTYRML) (57) or 50 μg/ml of total cellular antigen (infected or noninfected VeroE6 lysate). Following over-night incubation, the splenocytes were incubated for 2 h with 5 μg/ml brefeldin A (Biolegend) for intracellular trapping of cytokines and then stained with Zombie Aqua (1:200) in PBS for 15 min. Splenocytes were then stained for CD3 (4 μg/ml eFluor 450 a-mouse CD3 antibody; ThermoFisher Scientific) and CD8 (2 μg/ml APC/Cy7 a-mouse CD8a antibody; Biolegend) in PBS for 20 min before fixation in 2% paraformaldehyde (Sigma-Aldrich), and permeabilization and blocking in a mixture of 0.1% saponin and 1% normal mouse serum. Finally, splenocytes were stained intracellularly for TNFa (6.5 μg/ml PE anti-mouse TNFa, Biolegend) and IFN-γ (2 μg/ml APC a-mouse IFN-γ, Biolegend) prior to analysis on a flow cytometer (LSR Fortessa X-20, Becton Dickinson). Gating FSC-A/SSC-A excluded debris, SSC-H/SSC-W and FSC-H/FSC-W excluded doublet cells. The data were analysed using FlowJo 10 software (TreeStar). To determine the percentage of responding CD4^+^ or CD8^+^ T lymphocytes, the percentage of responders from samples stimulated with non-infected Vero E6 lysates were subtracted from the corresponding responses. The gating strategy for the analysis is depicted in Fig. S10B.

### Statistical analysis

Graph Pad Prism 7 (GraphPad Software, Inc.) was used for all statistical evaluations. Quantitative data were represented as mean ± standard deviation (SD) and obtained from at least three independent experiments. For ADCC assays, flow cytometry analysis and ELISpot assays data were represented as mean ± standard error of mean (SEM). Statistical significance was determined using survival analysis with log-rank (Mantel-Cox) test, one-way ANOVA analysis (neutralization titers and ELISA), two-way ANOVA analysis (ADCC), paired t-test (flow cytometry) and Wilcoxon matched pairs signed rank test (comparison of paired post-vaccinated and post-challenge samples). Correlation studies were performed using linear regression analysis with Pearson’s correlation coefficient. Values were considered statistically significantly different at p-values ≤ 0.05.

## SUPPLEMENTAL MATERIAL

Supplemental material for the article is added as Fig. S1–S10.

## ACKNOWLEDGEMENT

The authors thank Katrien Geerts and Sarah Debaveye for their excellent technical assistance, Jef Ceulemans and Jonas Piot for their contribution to assay development, and Robert Vrancken and Joeri Auwerx for assisting in the generation of recombinant JEV SA14-14-2.

## SUPPLEMENTAL MATERIAL

**FIG S1.**
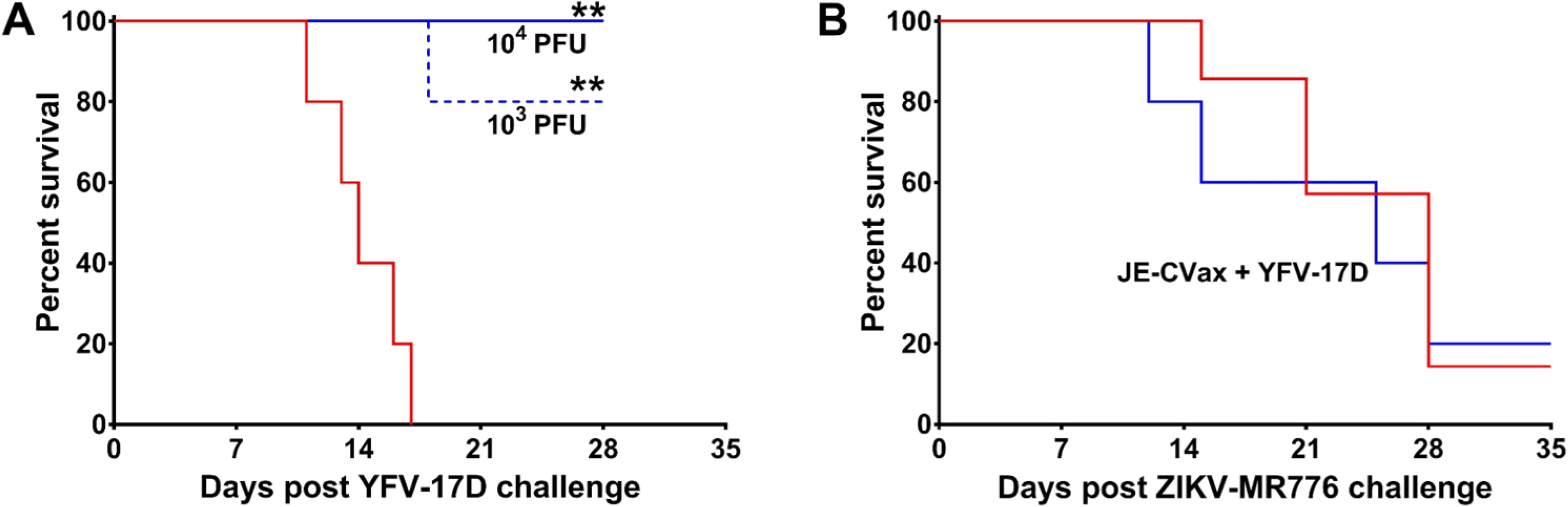
*In vivo* characterization of protective efficacy of JE-CVax vaccination against lethal YFV-17D and ZIKV-MR766 challenge in AG129 mice. **(A)** AG129 mice (n = 5) were vaccinated i.p. with 10^3-4^PFU of JE-CVax (blue) and 28 dpv, challenged i.p. with 10^3^ PFU of YFV-17D. **(B)** AG129 mice (n = 6) were first vaccinated i.p. with 10^4^ PFU JE-CVax and 28 dpv challenged i.p. with 10^3^ PFU YFV-17D (blue). 28 days post YFV-17D challenge animals were challenged a second time, yet with 10^4^ PFU ZIKV-MR766 and observed for mortality for the following 5 weeks. Age-matched non-vaccinated (red) animals were challenged with 10^3^ PFU YFV-17D [n = 5 **(A)**] or 10^4^ PFU ZIKA-MR766 [n = 7 **(B)**] as controls. Log-rank (Mantel-Cox) survival analysis test was performed for statistical significance. ** p-value ≤ 0.01 compared to non-vaccinated group.

**FIG S2.**
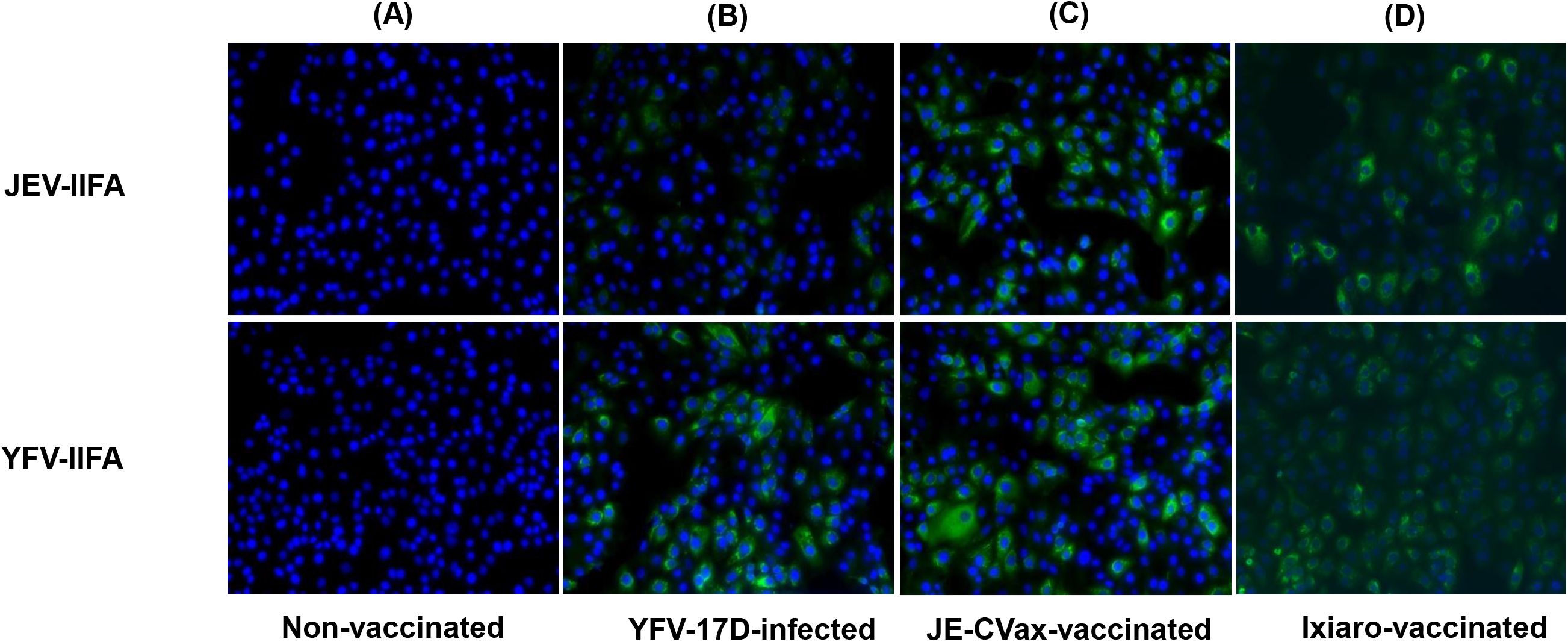
Detection of cross-reactive antibodies in sera of YFV-17D-infected and JE-CVax-vaccinated mouse serum against JEV and YFV. AG129 mice were pre-bled before either infection with 10^3^ PFU YFV-17D or vaccination with 10^4^ PFU JE-CVax/Ixiaro and bled again either at the onset of sickness (YFV-17D) or on 28 dpv (JE-CVax). Preserum **(A)**, serum of YFV-17D infected mice **(B)**, serum of JE-CVax-vaccinated mice **(B)** and serum of Ixiaro-vaccinated mice **(D)** were analyzed by both JEV (upper panel) and YFV (lower panel) indirect immunofluorescence assay (IIFA, Euroimmun^®^) at 20X magnification.

**FIG S3.**
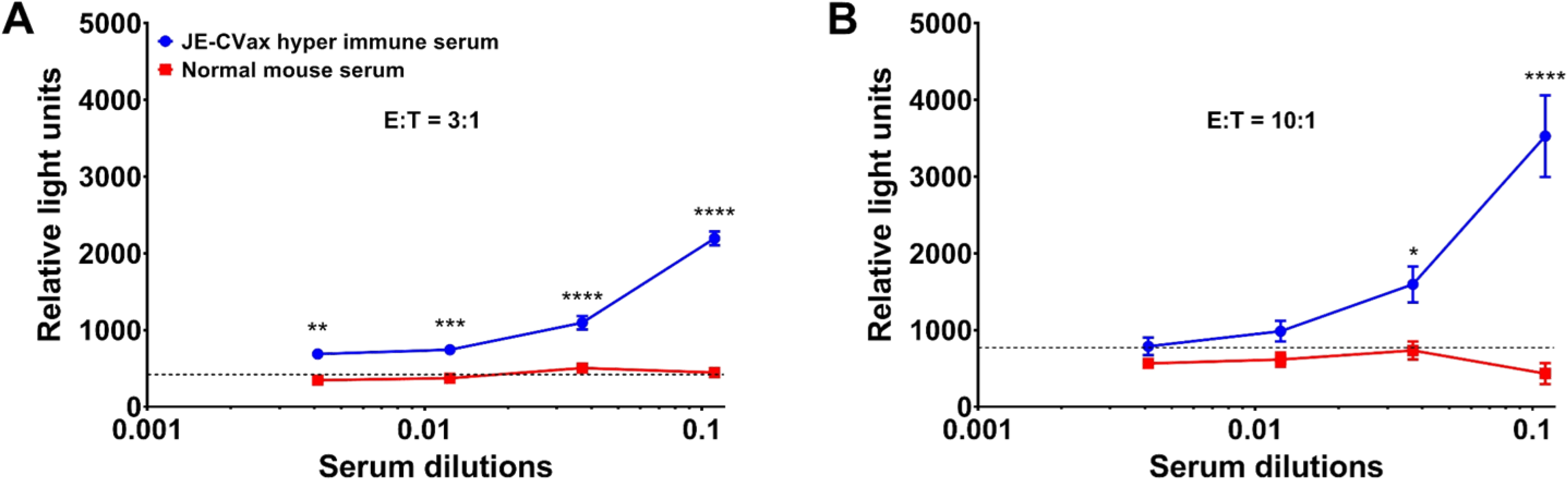
Effect of JE-CVax hyperimmune serum in antibody-dependent cellular cytotoxicity (ADCC). JE-CVax hyperimmune serum 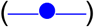 was tested for its ability to mediate ADCC activity in comparison to serum of non-vaccinated mice 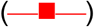; at 3:1 **(A)** and 10:1 **(B)** effector (E): target (T) ratios. Data presented are from experiments conducted twice, each in triplicate, and presented as mean ± SEM. The average of relative light unit signal plus three standard deviations from E:T in absence of hyperimmune serum was considered as the background signal (CC). Statistical significance was determined using two-way ANOVA analysis. *, **, ***, **** p-value ≤ 0.05, ≤ 0.01, ≤ 0.001 and ≤ 0.0001 compared to normal serum.

**FIG S4.**
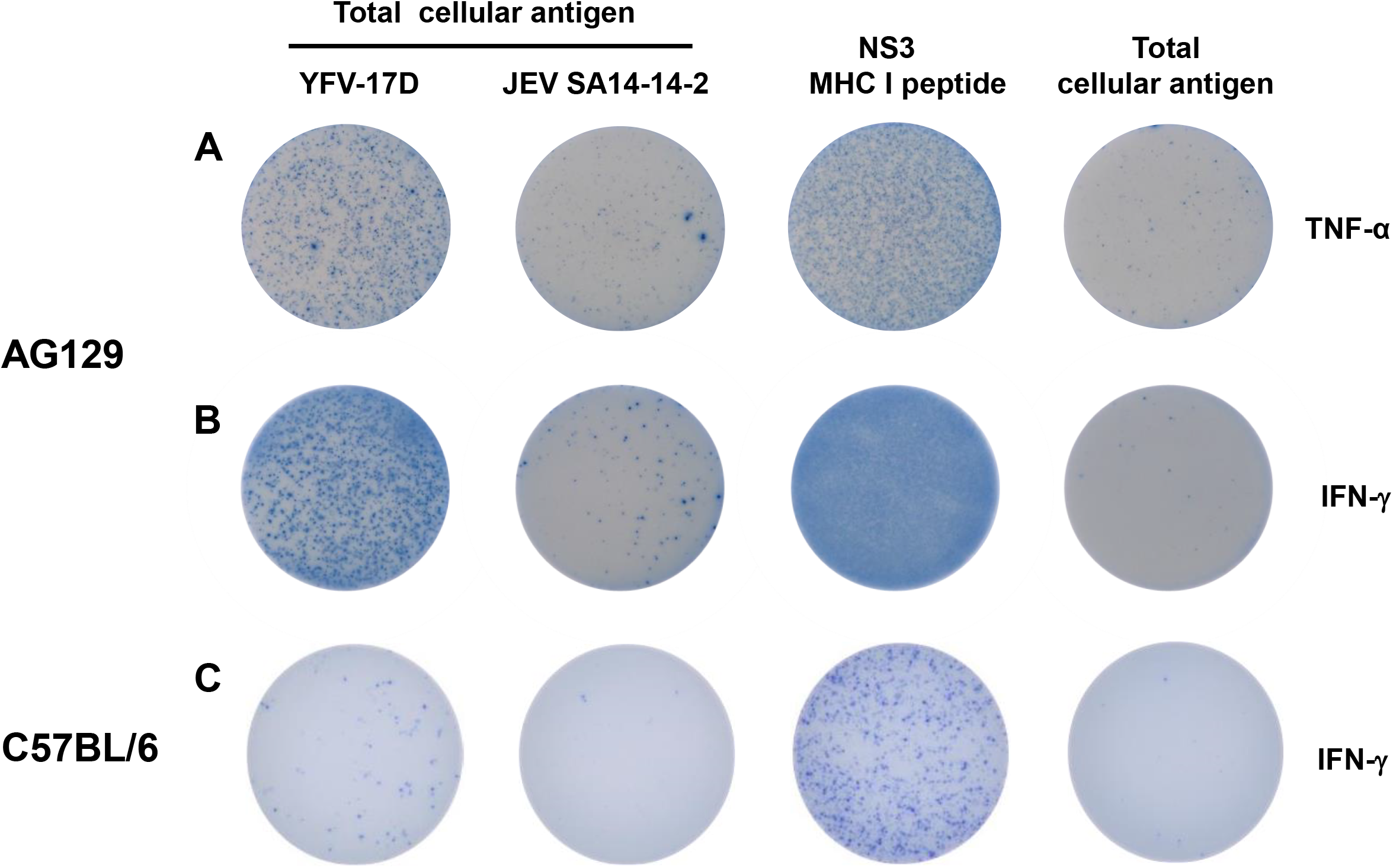
T cell responses directed against YFV or JEV antigens in ELISpot assay. Representative wells of ELISpot assays showing TNF-α **(A)** and IFN-γ **(B, C)** production by splenocytes of AG129 mice **(A, B)** and C57BL/6 mice **(C)**, at 18- and 4-weeks, respectively, post-vaccination with 10^4^ PFU JE-CVax, following 16 h *ex vivo* re-stimulation with a MHC class I restricted peptide derived from YFV NS3^32^, or the lysate of YFV-17D- or JEV SA14-14-2-infected Vero E6 cells. Stimulation using lysate of non-infected Vero E6 cells served as negative control.

**FIG S5.**
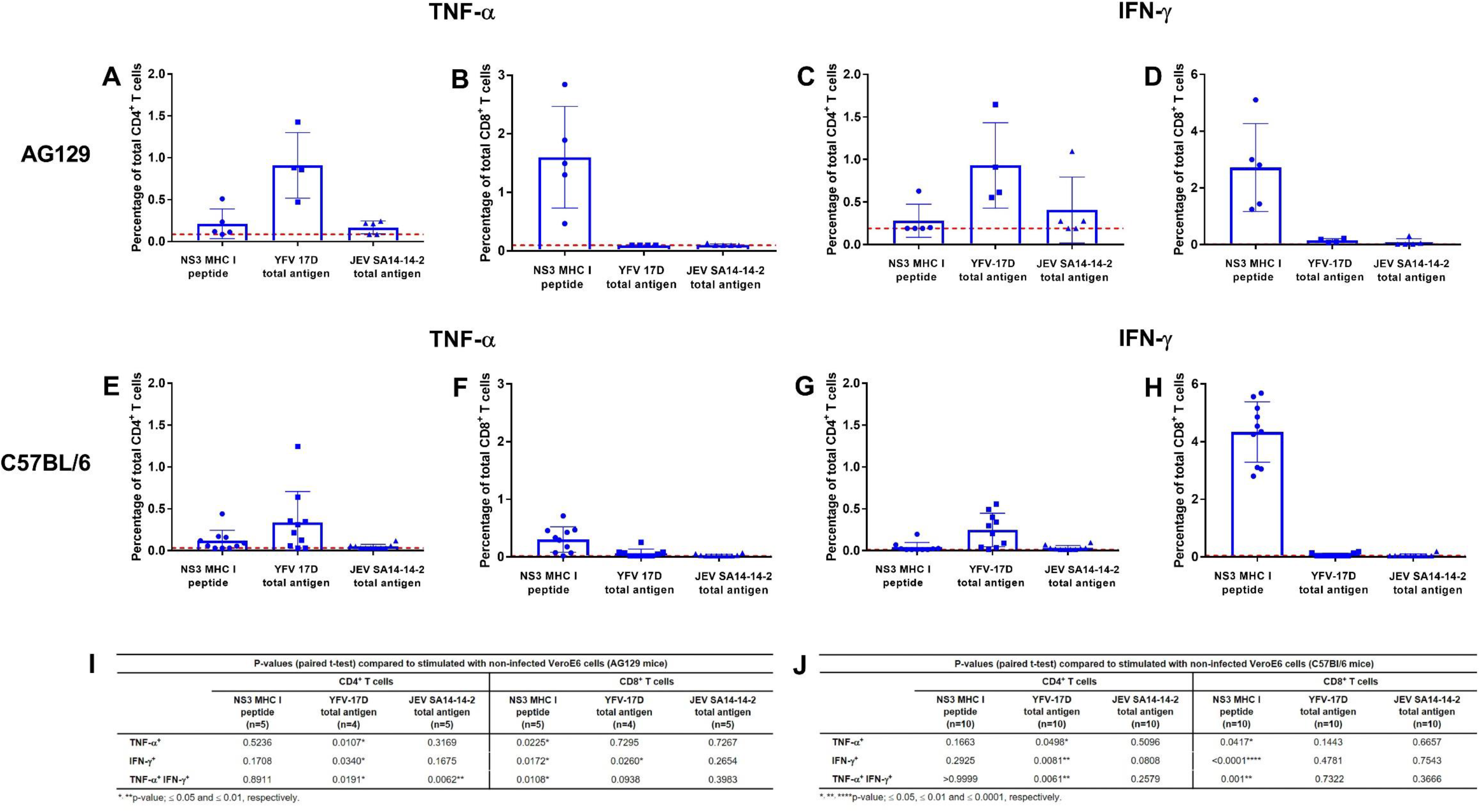
Detection of T cell responses in mice by intracellular staining for TNF-α and IFN-γ using flow cytometry. Flow cytometric analysis for intracellular TNF-α **(A, B, E, F)** and IFN-γ **(C, D, G, H)** production by **(A, C, E, G)** CD4^+^ or **(B, D, F, H)** CD8^+^ T cells from vaccinated AG129 mice (A-D) and C57BL/6 mice (EH) following stimulation with either NS3 MHC I peptide or cell lysate of YFV-17D or JEV SA14-14-2 infected VeroE6 cells. Percentage of total CD4^+^or CD8^+^ TNF-α or IFN-γ secreting T cells analyzed in flow cytometric analysis in AG129 mice (n = 5) and C57BL/6 mice (n = 10). **Table I and J** represents p-values between cytokine-secreting populations of antigen-versus non-infected VeroE6-stimulated samples. (Statistical significance; paired t-test) of flow cytometric analysis for splenocytes from AG129 and C57BL/6 mice, respectively. The data were compiled from two independent experiments and dotted lines represent average background in control samples collected from non-vaccinated animals.

**FIG S6.**
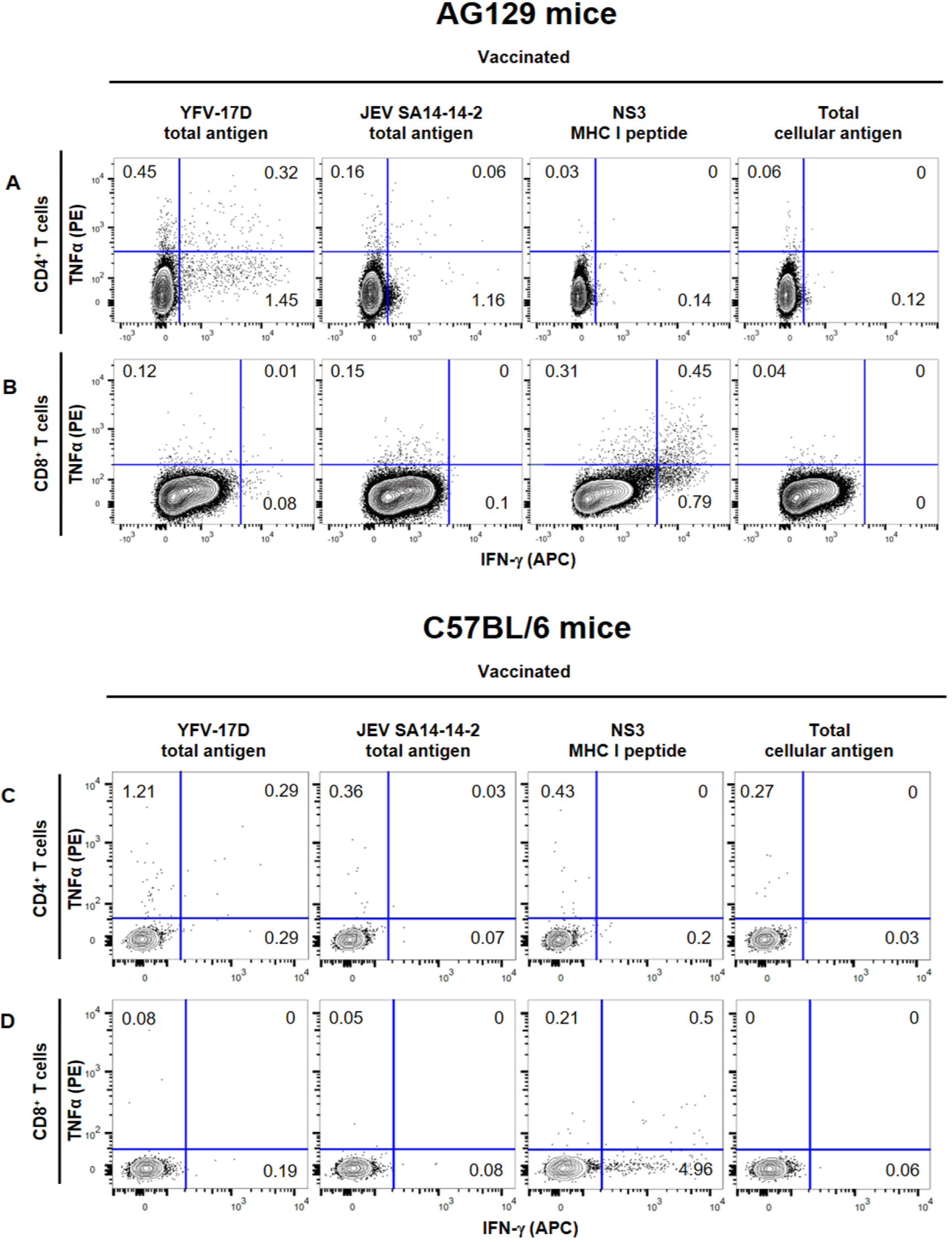
Detection of T cell responses through intracellular cytokine staining in AG129 and C57BL/6 mice. Representative depiction of flow cytometric analysis for intracellular TNF-α and IFN-γ production by **(A, C)** CD4^+^ (gated CD3^+^ CD8^−^) or **(B,D)** CD8^+^ T cells (gated CD3^+^ CD8^+^) from JE-CVax-vaccinated **(A, B)** AG129 mice, 18 weeks post-vaccination and **(C, D)** C57BL/6 mice, 4 weeks post-vaccination, following 16h *ex vivo* stimulation with cell lysate of YFV-17D or JEV SA14-14-2 infected Vero E6 cells, a MHC I restricted NS3 peptide, or uninfected Vero E6 cells.

**FIG S7.**
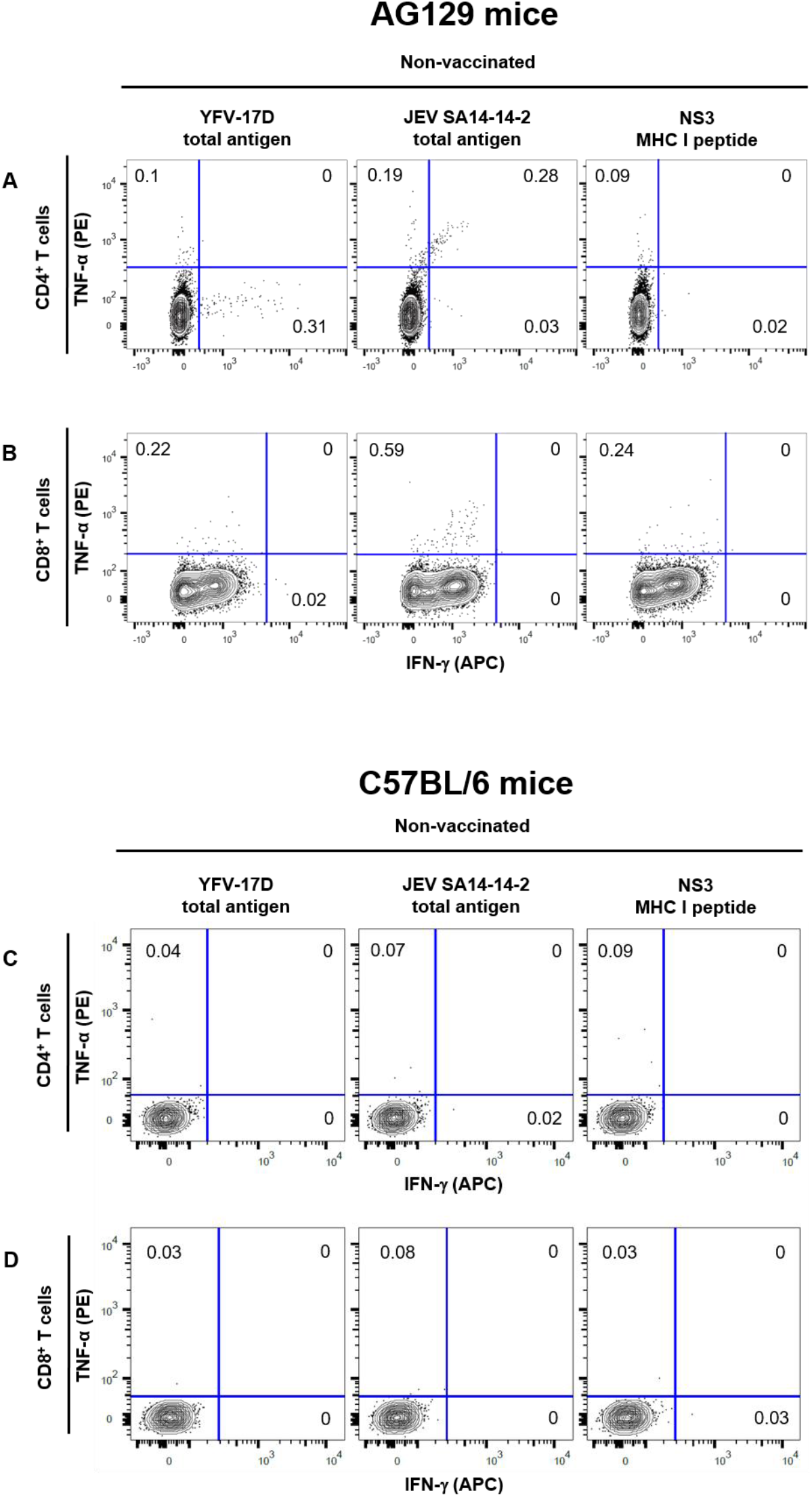
Detection of T cell responses through intracellular cytokine staining in non-vaccinated baseline AG 129 and C57BL/6 mice controls. Representative depiction of flow cytometric analysis for intracellular TNF-α and IFN-γ production by **(A, C)** CD4^+^ (gated CD3^+^ CD8^−^) or **(B, D)** CD8^+^ T cells (gated CD3^+^ CD8^+^) from non-vaccinated AG129 mice following 16h ex *vivo* stimulation with cell lysates of YFV-17D or JEV SA14-14-2 infected Vero E6 cells, or a MHC I restricted YFV NS3 peptide.

**FIG S8.**
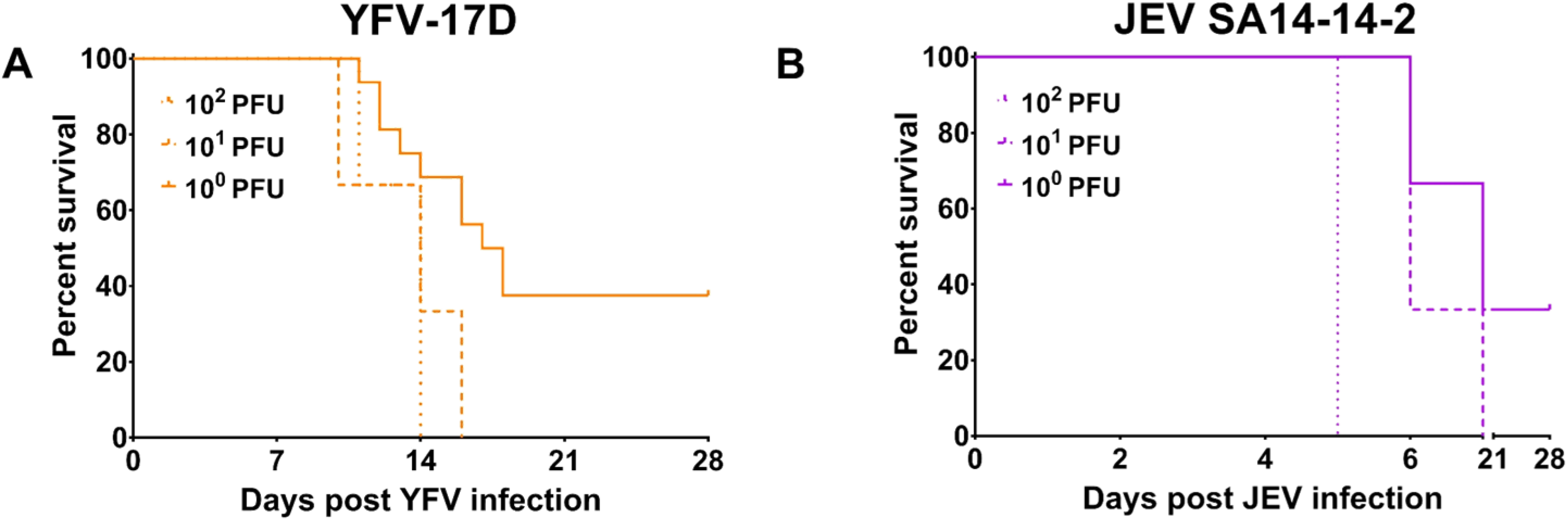
*In vivo* infectivity of YFV-17D and JEV SA14-14-2 in AG129 mice. AG129 mice were inoculated via the i.p. route with different doses of **(A)** YFV-17D [10^0^ (—; n = 16), 10^1^ (- -; n = 3) or 10^2^ PFU (….; n = 3)], or **(B)** JEV SA14-14-2 [10^0^ (—), 10^1^ (- -) or 10^2^ PFU (….), n =3]. Animals were monitored over a period of five weeks and were euthanized when humane end-points were reached.

**FIG S9.**
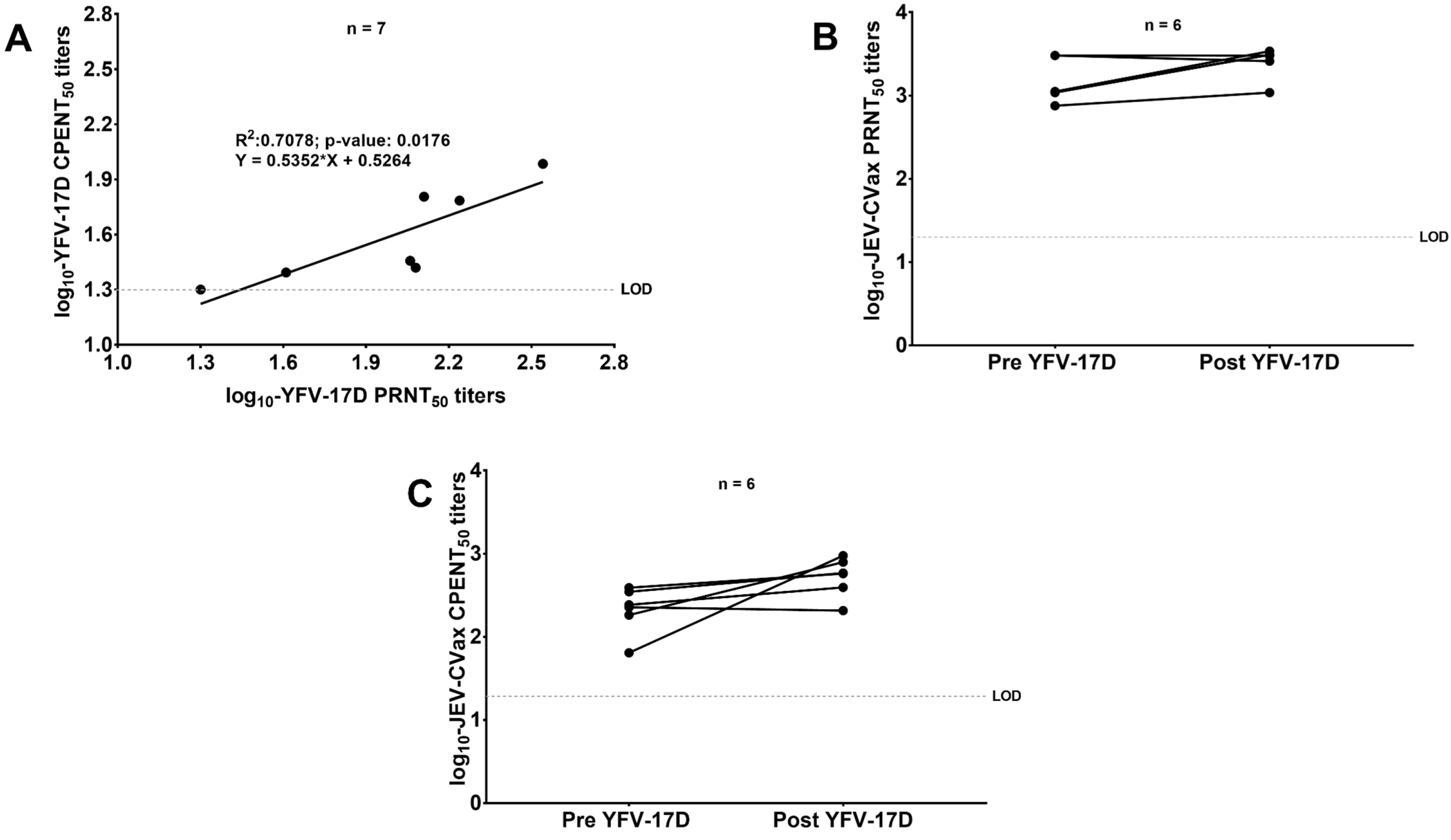
Correlation of nAb titers determined as log_10_PRNT_50_ and log_10_CPENT_50_ in matched serum samples of JE-CVax-vaccinated and/or YFV-1 ZD-challenged mice. AG129 mice (n = 7) were vaccinated with 10^4^ PFU of JE-CVax and 28 dpv challenged with 10^3^ PFU YFV-17D. Serum was harvested and neutralization assays i.e. CPENT and PRNT were performed as described in Materials and Methods. Data presented show a good correlation (Pearson correlation: R^2^ = 0.071; p = 0.02) between log_10_-YFV PRNT_50_ and log_10_-YFV CPENT_50_ of matched samples **(A)**. There was no marked increase in the log_10_-JE-CVax PRNT_50_ and log_10_-JE-CVax CPENT_50_ titers (p-value; 0.156 and 0.062, respectively) when comparing matched serum samples from before and after challenge with YFV-17D using Wilcoxon matched pairs signed rank test **(B, C)**. Limit of detection for either assay was log_10_20 i.e. 1.3.

**FIG S10.**
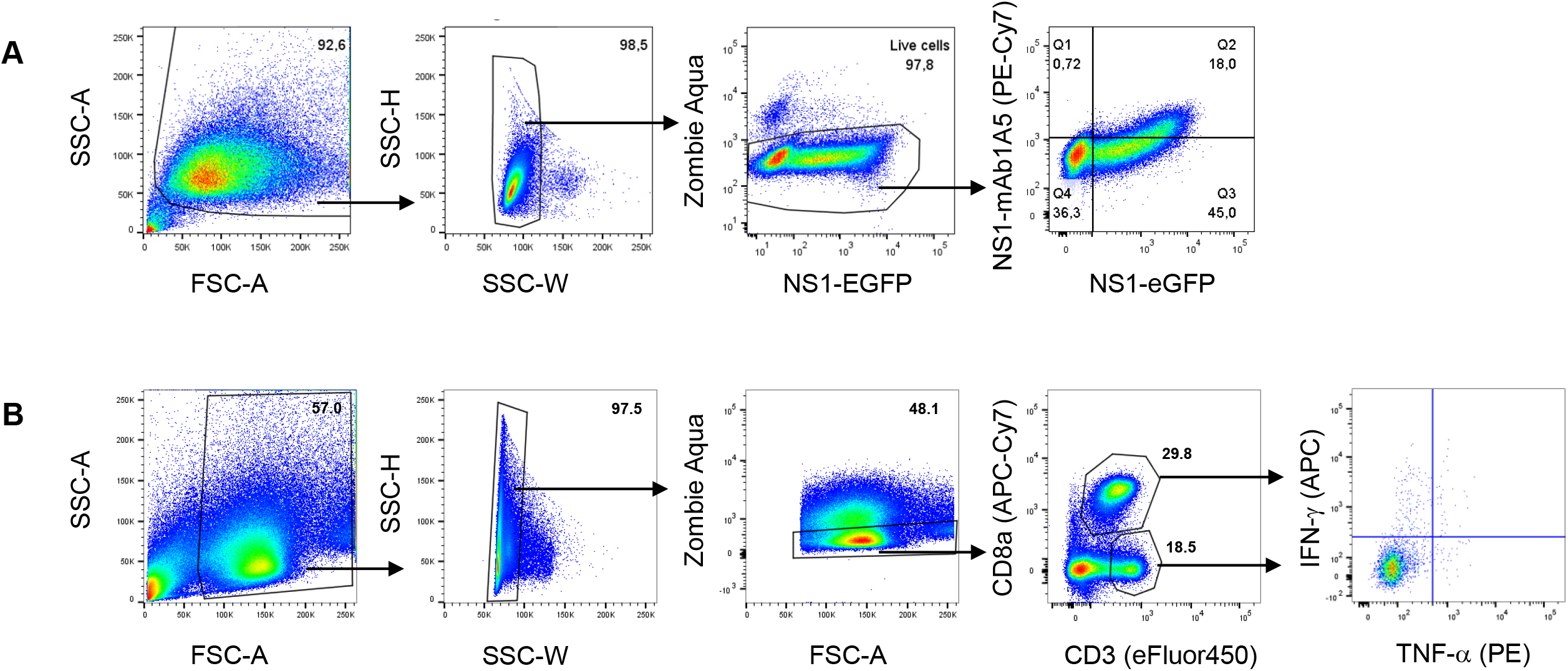
Gating strategy for flow cytometry analysis. **(A, B)** Exclusion of debris was achieved by gating out the FSC-low population in a FSC-A vs SSC-A plot. Then, only single cells were retained by elimination of the high SSC-W population, in a SSC-W vs SSC-H plot. In a subsequent step, live cells were selected by gating out the Zombie Aqua-positive population as shown here in an NS1-eGFP/FSC-A vs ZA plot. Finally for the detection of anti-NS1 antibodies **(A)** cells were gated based on positivity for NS1-eGFP and positivity/negativity of a-NS1 Ab PE-Cy7. **(B)** For intracellular cytokine staining, based on positivity for CD3e (eFluor450-conjugated) and negativity (CD4) or positivity for CD8a (APC/Cy7-conjugated), CD4^+^ and CD8^+^ T cell populations were defined as CD3^+^CD8^−^ and CD3^+^CD8^+^ populations, respectively. Finally, cells were gated based on positivity/negativity for IFN-γ and positivity/negativity of TNF-α. Samples from non-vaccinated mice were used to set the boundaries that define cells positive and negative for intracellular markers.

## REFERENCES

1. Ishikawa T, Yamanaka A, Konishi E. 2014. A review of successful flavivirus vaccines and the problems with those flaviviruses for which vaccines are not yet available. Vaccine 32:1326–37.

2. Petersen LR, Jamieson DJ, Powers AM, Honein MA. 2016. Zika Virus. N Engl J Med 374:1552–63.

3. Wilder-Smith A, Monath TP. 2017. Responding to the threat of urban yellow fever outbreaks. Lancet Infect Dis 17:248–250.

4. Simon-Loriere E, Faye O, Prot M, Casademont I, Fall G, Fernandez-Garcia MD, Diagne MM, Kipela JM, Fall IS, Holmes EC, Sakuntabhai A, Sall AA. 2017. Autochthonous Japanese Encephalitis with Yellow Fever Coinfection in Africa. N Engl J Med 376:1483–1485.

5. Who. 2015. Vaccines and vaccination against yellow fever: WHO Position Paper, June 2013--recommendations. Vaccine 33:76–7.

6. Barrett AD. 2016. Yellow Fever in Angola and Beyond--The Problem of Vaccine Supply and Demand. N Engl J Med 375:301–3.

7. Gershman MD, Angelo KM, Ritchey J, Greenberg DP, Muhammad RD, Brunette G, Cetron MS, Sotir MJ. 2017. Addressing a Yellow Fever Vaccine Shortage – United States, 2016-2017. MMWR Morb Mortal Wkly Rep 66:457–459.

8. Wasserman S, Tambyah PA, Lim PL. 2016. Yellow fever cases in Asia: primed for an epidemic. Int J Infect Dis 48:98–103.

9. Chen Z, Liu L, Lv Y, Zhang W, Li J, Zhang Y, Di T, Zhang S, Liu J, Qu J, Hua W, Li C, Wang P, Zhang Q, Xu Y, Jiang R, Wang Q, Chen L, Wang S, Pang X, Liang M, Ma X, Li X, Zhang F, Li D. 2016. A fatal yellow fever virus infection in China: description and lessons. Emerg Microbes Infect 5:e69.

10. Li X, Ma SJ, Liu X, Jiang LN, Zhou JH, Xiong YQ, Ding H, Chen Q. 2014. Immunogenicity and safety of currently available Japanese encephalitis vaccines: a systematic review. Hum Vaccin Immunother 10:3579–93.

11. Monath TP, Seligman SJ, Robertson JS, Guy B, Hayes EB, Condit RC, Excler JL, Mac LM, Carbery B, Chen RT, BCVVVSWG. 2015. Live virus vaccines based on a yellow fever vaccine backbone: standardized template with key considerations for a risk/benefit assessment. Vaccine 33:62–72.

12. Dans AL, Dans LF, Lansang MAD, Silvestre MAA, Guyatt GH. 2018. Controversy and debate on dengue vaccine series-paper 1: review of a licensed dengue vaccine: inappropriate subgroup analyses and selective reporting may cause harm in mass vaccination programs. J Clin Epidemiol 95:137–139.

13. Aguiar M, Stollenwerk N. 2018. Dengvaxia Efficacy Dependency on Serostatus: A Closer Look at More Recent Data. Clin Infect Dis 66:641–642.

14. Wilder-Smith A. 2018. Four-year safety follow-up of the tetravalent dengue vaccine CYD-TDV. Clin Microbiol Infect 24:680–681.

15. Pierson TC, Fremont DH, Kuhn RJ, Diamond MS. 2008. Structural insights into the mechanisms of antibody-mediated neutralization of flavivirus infection: implications for vaccine development. Cell Host Microbe 4:229–38.

16. Watson AM, Lam LK, Klimstra WB, Ryman KD. 2016. The 17D-204 Vaccine Strain-Induced Protection against Virulent Yellow Fever Virus Is Mediated by Humoral Immunity and CD4+ but not CD8+ T Cells. PLoS Pathog 12:e1005786.

17. Putnak JR, Schlesinger JJ. 1990. Protection of mice against yellow fever virus encephalitis by immunization with a vaccinia virus recombinant encoding the yellow fever virus non-structural proteins, NS1, NS2a and NS2b. J Gen Virol 71 (Pt 8):1697–702.

18. Schlesinger JJ, Foltzer M, Chapman S. 1993. The Fc portion of antibody to yellow fever virus NS1 is a determinant of protection against YF encephalitis in mice. Virology 192:132–41.

19. Schlesinger JJ, Brandriss MW, Cropp CB, Monath TP. 1986. Protection against yellow fever in monkeys by immunization with yellow fever virus nonstructural protein NS1. J Virol 60:1153–5.

20. Rastogi M, Sharma N, Singh SK. 2016. Flavivirus NS1: a multifaceted enigmatic viral protein. Virol J 13:131.

21. Chung KM, Thompson BS, Fremont DH, Diamond MS. 2007. Antibody recognition of cell surface-associated NS1 triggers Fc-gamma receptor-mediated phagocytosis and clearance of West Nile Virus-infected cells. J Virol 81:9551–5.

22. Bailey MJ, Broecker F, Duehr J, Arumemi F, Krammer F, Palese P, Tan GS. 2019. Antibodies Elicited by an NS1-Based Vaccine Protect Mice against Zika Virus. MBio 10.

23. Bassi MR, Larsen MA, Kongsgaard M, Rasmussen M, Buus S, Stryhn A, Thomsen AR, Christensen JP. 2016. Vaccination with Replication Deficient Adenovectors Encoding YF-17D Antigens Induces Long-Lasting Protection from Severe Yellow Fever Virus Infection in Mice. PLoS Negl Trop Dis 10:e0004464.

24. Desprès P, Dietrich J, Girard M, Bouloy M. 1991. Recombinant baculoviruses expressing yellow fever virus E and NS1 proteins elicit protective immunity in mice. J Gen Virol 72 (Pt 11):2811–6.

25. Chan KR, Wang X, Saron WAA, Gan ES, Tan HC, Mok DZL, Zhang SL, Lee YH, Liang C, Wijaya L, Ghosh S, Cheung YB, Tannenbaum SR, Abraham SN, St John AL, Low JGH, Ooi EE. 2016. Cross-reactive antibodies enhance live attenuated virus infection for increased immunogenicity. Nat Microbiol 1:16164.

26. Li J, Gao N, Fan D, Chen H, Sheng Z, Fu S, Liang G, An J. 2016. Cross-protection induced by Japanese encephalitis vaccines against different genotypes of Dengue viruses in mice. Sci Rep 6:19953.

27. Schlesinger JJ, Brandriss MW, Monath TP. 1983. Monoclonal antibodies distinguish between wild and vaccine strains of yellow fever virus by neutralization, hemagglutination inhibition, and immune precipitation of the virus envelope protein. Virology 125:8–17.

28. Kum DB, Mishra N, Boudewijns R, Gladwyn-Ng I, Alfano C, Ma J, Schmid MA, Marques RE, Schols D, Kaptein S, Nguyen L, Neyts J, Dallmeier K. 2018. A yellow fever-Zika chimeric virus vaccine candidate protects against Zika infection and congenital malformations in mice. NPJ Vaccines 3:56.

29. Cheng ZJ, Garvin D, Paguio A, Moravec R, Engel L, Fan F, Surowy T. 2014. Development of a robust reporter-based ADCC assay with frozen, thaw-and-use cells to measure Fc effector function of therapeutic antibodies. J Immunol Methods 414:69–81.

30. Akondy RS, Monson ND, Miller JD, Edupuganti S, Teuwen D, Wu H, Quyyumi F, Garg S, Altman JD, Del Rio C, Keyserling HL, Ploss A, Rice CM, Orenstein WA, Mulligan MJ, Ahmed R. 2009. The yellow fever virus vaccine induces a broad and polyfunctional human memory CD8+ T cell response. J Immunol 183:7919–30.

31. Li XF, Deng YQ, Yang HQ, Zhao H, Jiang T, Yu XD, Li SH, Ye Q, Zhu SY, Wang HJ, Zhang Y, Ma J, Yu YX, Liu ZY, Li YH, Qin ED, Shi PY, Qin CF. 2013. A chimeric dengue virus vaccine using Japanese encephalitis virus vaccine strain SA14-14-2 as backbone is immunogenic and protective against either parental virus in mice and nonhuman primates. J Virol 87:13694–705.

32. Bassi MR, Kongsgaard M, Steffensen MA, Fenger C, Rasmussen M, Skjødt K, Finsen B, Stryhn A, Buus S, Christensen JP, Thomsen AR. 2015. CD8+ T cells complement antibodies in protecting against yellow fever virus. J Immunol 194:1141–53.

33. Calvert AE, Dixon KL, Delorey MJ, Blair CD, Roehrig JT. 2014. Development of a small animal peripheral challenge model of Japanese encephalitis virus using interferon deficient AG129 mice and the SA14-14-2 vaccine virus strain. Vaccine 32:258–64.

34. Thibodeaux BA, Garbino NC, Liss NM, Piper J, Blair CD, Roehrig JT. 2012. A small animal peripheral challenge model of yellow fever using interferon-receptor deficient mice and the 17D-204 vaccine strain. Vaccine 30:3180–7.

35. Meier KC, Gardner CL, Khoretonenko MV, Klimstra WB, Ryman KD. 2009. A mouse model for studying viscerotropic disease caused by yellow fever virus infection. PLoS Pathog 5:e1000614.

36. Yang J, Yang H, Li Z, Lin H, Zhao Y, Wang W, Tan S, Zeng X, Li Y. 2017. The chimeric Japanese encephalitis/Dengue 2 virus protects mice from challenge by both dengue virus and JEV virulent virus. Protein Cell 8:225–229.

37. Zellweger RM, Tang WW, Eddy WE, King K, Sanchez MC, Shresta S. 2015. CD8+ T Cells Can Mediate Short-Term Protection against Heterotypic Dengue Virus Reinfection in Mice. J Virol 89:6494–505.

38. Xu X, Vaughan K, Weiskopf D, Grifoni A, Diamond MS, Sette A, Peters B. 2016. Identifying Candidate Targets of Immune Responses in Zika Virus Based on Homology to Epitopes in Other Flavivirus Species. PLoS Curr 8.

39. Nasveld PE, Marjason J, Bennett S, Aaskov J, Elliott S, McCarthy K, Kanesa-Thasan N, Feroldi E, Reid M. 2010. Concomitant or sequential administration of live attenuated Japanese encephalitis chimeric virus vaccine and yellow fever 17D vaccine: randomized double-blind phase II evaluation of safety and immunogenicity. Hum Vaccin 6:906–14.

40. Chambers TJ, Nestorowicz A, Mason PW, Rice CM. 1999. Yellow fever/Japanese encephalitis chimeric viruses: construction and biological properties. J Virol 73:3095–101.

41. Monath TP, Soike K, Levenbook I, Zhang ZX, Arroyo J, Delagrave S, Myers G, Barrett AD, Shope RE, Ratterree M, Chambers TJ, Guirakhoo F. 1999. Recombinant, chimaeric live, attenuated vaccine (ChimeriVax) incorporating the envelope genes of Japanese encephalitis (SA14-14-2) virus and the capsid and nonstructural genes of yellow fever (17D) virus is safe, immunogenic and protective in non-human primates. Vaccine 17:1869–82.

42. Stiasny K, Kiermayr S, Holzmann H, Heinz FX. 2006. Cryptic properties of a cluster of dominant flavivirus cross-reactive antigenic sites. J Virol 80:9557–68.

43. Schlesinger JJ, Brandriss MW, Putnak JR, Walsh EE. 1990. Cell surface expression of yellow fever virus non-structural glycoprotein NS1: consequences of interaction with antibody. J Gen Virol 71 (Pt 3):593–9.

44. Gould EA, Buckley A, Barrett AD, Cammack N. 1986. Neutralizing (54K) and non-neutralizing (54K and 48K) monoclonal antibodies against structural and non-structural yellow fever virus proteins confer immunity in mice. J Gen Virol 67 (Pt 3):591–5.

45. James EA, LaFond RE, Gates TJ, Mai DT, Malhotra U, Kwok WW. 2013. Yellow fever vaccination elicits broad functional CD4+ T cell responses that recognize structural and nonstructural proteins. J Virol 87:12794–804.

46. Barrett AD, Gould EA. 1986. Comparison of neurovirulence of different strains of yellow fever virus in mice. J Gen Virol 67 (Pt 4):631–7.

47. Perelygin AA, Scherbik SV, Zhulin IB, Stockman BM, Li Y, Brinton MA. 2002. Positional cloning of the murine flavivirus resistance gene. Proc Natl Acad Sci U S A 99:9322–7.

48. Theiler M, Smith HH. 1937. THE USE OF YELLOW FEVER VIRUS MODIFIED BY IN VITRO CULTIVATION FOR HUMAN IMMUNIZATION. J Exp Med 65:787–800.

49. Monath TP, Guirakhoo F, Nichols R, Yoksan S, Schrader R, Murphy C, Blum P, Woodward S, McCarthy K, Mathis D, Johnson C, Bedford P. 2003. Chimeric live, attenuated vaccine against Japanese encephalitis (ChimeriVax-JE): phase 2 clinical trials for safety and immunogenicity, effect of vaccine dose and schedule, and memory response to challenge with inactivated Japanese encephalitis antigen. J Infect Dis 188:1213–30.

50. Guirakhoo F, Kitchener S, Morrison D, Forrat R, McCarthy K, Nichols R, Yoksan S, Duan X, Ermak TH, Kanesa-Thasan N, Bedford P, Lang J, Quentin-Millet MJ, Monath TP. 2006. Live attenuated chimeric yellow fever dengue type 2 (ChimeriVax-DEN2) vaccine: Phase I clinical trial for safety and immunogenicity: effect of yellow fever pre-immunity in induction of cross neutralizing antibody responses to all 4 dengue serotypes. Hum Vaccin 2:60–7.

51. Xie X, Yang Y, Muruato AE, Zou J, Shan C, Nunes BT, Medeiros DB, Vasconcelos PF, Weaver SC, Rossi SL, Shi PY. 2017. Understanding Zika Virus Stability and Developing a Chimeric Vaccine through Functional Analysis. MBio 8.

52. Arroyo J, Guirakhoo F, Fenner S, Zhang ZX, Monath TP, Chambers TJ. 2001. Molecular basis for attenuation of neurovirulence of a yellow fever Virus/Japanese encephalitis virus chimera vaccine (ChimeriVax-JE). J Virol 75:934–42.

53. Zmurko J, Marques RE, Schols D, Verbeken E, Kaptein SJ, Neyts J. 2016. The Viral Polymerase Inhibitor 7-Deaza-2’-C-Methyladenosine Is a Potent Inhibitor of In Vitro Zika Virus Replication and Delays Disease Progression in a Robust Mouse Infection Model. PLoS Negl Trop Dis 10:e0004695.

54. Bredenbeek PJ, Kooi EA, Lindenbach B, Huijkman N, Rice CM, Spaan WJ. 2003. A stable full-length yellow fever virus cDNA clone and the role of conserved RNA elements in flavivirus replication. J Gen Virol 84:1261–8.

55. Demidenko AA, Blattman JN, Blattman NN, Greenberg PD, Nibert ML. 2013. Engineering recombinant reoviruses with tandem repeats and a tetravirus 2A-like element for exogenous polypeptide expression. Proc Natl Acad Sci U S A 110:E1867–76.

56. Avirutnan P, Fuchs A, Hauhart RE, Somnuke P, Youn S, Diamond MS, Atkinson JP. 2010. Antagonism of the complement component C4 by flavivirus nonstructural protein NS1. J Exp Med 207:793–806.

57. Barba-Spaeth G, Longman RS, Albert ML, Rice CM. 2005. Live attenuated yellow fever 17D infects human DCs and allows for presentation of endogenous and recombinant T cell epitopes. J Exp Med 202:1179–84.

58. Li XF, Li XD, Deng CL, Dong HL, Zhang QY, Ye Q, Ye HQ, Huang XY, Deng YQ, Zhang B, Qin CF. 2017. Visualization of a neurotropic flavivirus infection in mouse reveals unique viscerotropism controlled by host type I interferon signaling. Theranostics 7:912–925.

59. Aguirre S, Maestre AM, Pagni S, Patel JR, Savage T, Gutman D, Maringer K, Bernal-Rubio D, Shabman RS, Simon V, Rodriguez-Madoz JR, Mulder LC, Barber GN, Fernandez-Sesma A. 2012. DENV inhibits type I IFN production in infected cells by cleaving human STING. PLoS Pathog 8:e1002934.

60. Ashour J, Morrison J, Laurent-Rolle M, Belicha-Villanueva A, Plumlee CR, Bernal-Rubio D, Williams KL, Harris E, Fernandez-Sesma A, Schindler C, Garcia-Sastre A. 2010. Mouse STAT2 restricts early dengue virus replication. Cell Host Microbe 8:410–21.

61. Grant A, Ponia SS, Tripathi S, Balasubramaniam V, Miorin L, Sourisseau M, Schwarz MC, Sanchez-Seco MP, Evans MJ, Best SM, Garcia-Sastre A. 2016. Zika Virus Targets Human STAT2 to Inhibit Type I Interferon Signaling. Cell Host Microbe 19:882–90.

62. Erickson AK, Pfeiffer JK. 2015. Spectrum of disease outcomes in mice infected with YFV-17D. J Gen Virol 96:1328–39.

63. Laurent-Rolle M, Boer EF, Lubick KJ, Wolfinbarger JB, Carmody AB, Rockx B, Liu W, Ashour J, Shupert WL, Holbrook MR, Barrett AD, Mason PW, Bloom ME, García-Sastre A, Khromykh AA, Best SM. 2010. The NS5 protein of the virulent West Nile virus NY99 strain is a potent antagonist of type I interferon-mediated JAK-STAT signaling. J Virol 84:3503–15.

64. Muñoz-Jordán JL, Laurent-Rolle M, Ashour J, Martínez-Sobrido L, Ashok M, Lipkin WI, García-Sastre A. 2005. Inhibition of alpha/beta interferon signaling by the NS4B protein of flaviviruses. J Virol 79:8004–13.

65. Julander JG. 2016. Animal models of yellow fever and their application in clinical research. Curr Opin Virol 18:64–9.

66. Lam LKM, Watson AM, Ryman KD, Klimstra WB. 2018. Gamma-interferon exerts a critical early restriction on replication and dissemination of yellow fever virus vaccine strain 17D-204. NPJ Vaccines 3:5.

67. Huang S, Hendriks W, Althage A, Hemmi S, Bluethmann H, Kamijo R, Vilcek J, Zinkernagel RM, Aguet M. 1993. Immune response in mice that lack the interferon-gamma receptor. Science 259:1742–5.

68. Kruisbeek AM. 2001. In vivo depletion of CD4-and CD8-specific T cells. Curr Protoc Immunol Chapter 4:Unit 4.1.

69. Johnson BW, Kosoy O, Hunsperger E, Beltran M, Delorey M, Guirakhoo F, Monath T. 2009. Evaluation of chimeric Japanese encephalitis and dengue viruses for use in diagnostic plaque reduction neutralization tests. Clin Vaccine Immunol 16:1052–9.

70. Reed LJ, Muench H. 1938. A SIMPLE METHOD OF ESTIMATING FIFTY PER CENT ENDPOINTS12. American Journal of Epidemiology 27:493–497.

71. Jochmans D, Leyssen P, Neyts J. 2012. A novel method for high-throughput screening to quantify antiviral activity against viruses that induce limited CPE. J Virol Methods 183:176–9.

72. Chao DY, Galula JU, Shen WF, Davis BS, Chang GJ. 2015. Nonstructural protein 1-specific immunoglobulin M and G antibody capture enzyme-linked immunosorbent assays in diagnosis of flaviviral infections in humans. J Clin Microbiol 53:557–66.

73. Falgout B, Bray M, Schlesinger JJ, Lai CJ. 1990. Immunization of mice with recombinant vaccinia virus expressing authentic dengue virus nonstructural protein NS1 protects against lethal dengue virus encephalitis. J Virol 64:4356–63.

